# Cell-nanoplastics association impacts cell proliferation and motility

**DOI:** 10.64898/2026.04.03.716369

**Authors:** Qin Ni, Jingyao Ma, Jinyu Fu, LaDaisha Thompson, Zhuoxu Ge, Dean Sharif, Yining Zhu, Hai-Quan Mao, Jude M. Phillip, Sean X. Sun

## Abstract

**Detection of micro- and nanoplastics (MNPs) in human tissues has raised growing concern about their biological effects on tissue and cell function. While previous studies have examined MNP-cell interaction, most focused on limited cell and plastic types. Here, we present a comprehensive, quantitative investigation into how different types of nanoplastics (NPs) associate with and affect diverse cell types under physiologically relevant conditions. Using microfluidic-calibrated fluorescence microscopy, we quantify NP accumulation in cells in vitro and match cellular NP concentrations to levels reported in human tissues. While cell-associated NPs could be gradually released in vitro, they persist in vivo for over one month without detectable reduction in a mouse model. We discover that NP exposure at these levels broadly impairs cell proliferation across epithelial, endothelial, fibroblast, and immune cells, with cell type-dependent sensitivity. NP exposure also reduces motility in T cells and fibroblasts, with more complex effects observed in macrophages. Mechanistically, NP-cell association and trans-epithelial transport involved not only classical endocytic regulators but also pathways related to ion and water transport. Notably, NP association and release were highly sensitive to the extracellular fluid environment within the physiological range. By testing inhibitors of these pathways, we identified molecules that reduce NP-cell association and promote release. We further compared common NPs found in human samples and widely used in research: polystyrene (PS), polyethylene (PE), and polypropylene (PP). Although these NPs similarly impaired proliferation and motility, they showed markedly different cellular association and release dynamics. These findings reveal the impact of NPs on tissue cell functions and uncover novel regulatory pathways, establishing a quantitative framework for studying NP-cell interactions in biologically relevant conditions.**

## Introduction

Exposure to microplastics (MPs) and nanoplastics (NPs) has emerged as a global health concern. MNPs are typically defined as plastic particles ranging from nanometers to tens of microns in size. While the sources of MNPs vary, ingestion through food and water is believed to be the primary route of exposure, with some studies also reporting uptake via the airway and direct dermal absorption (*1, 2*). Recent findings have revealed substantial MP accumulation in various human organs (*3–7*). In particular, MNP levels in decedent human brain tissue have been quantified at up to 0.2% of the wet tissue mass fraction (*3*), although recent work has called this into question (*8, 9*). Moreover, the concentration of MNPs in human samples has increased significantly over recent decades, with significantly higher accumulation of MNPs found in decedents with diagnosed dementia (*3*). Animal studies have shown that MNPs can rapidly translocate through the gastrointestinal tract, circulate through the bloodstream, and even cross the blood-brain barrier (*10–12*). Given that MNPs are often larger than cell-cell junctional space, their transport across tissue barriers is expected to involve active mechanisms such as endocytosis and exocytosis (*13*).

Whether and how MNPs affect fundamental tissue cell functions remains a central but unresolved question. Although largely chemically inert, MNPs act as rigid foreign bodies once internalized, and are likely to interfere with cytoskeletal organization and remodeling. This is particularly relevant for NPs of ∼100 nm, a size comparable to the mesh size of the actin cytoskeleton (*14*). At this critical length scale, NPs have complicated impacts on actin network mechanics and dynamics (*15*), potentially impairing intracellular trafficking that disrupts cell proliferation and migration. Although some studies have explored the impact of MNPs on cell proliferation and migration (*16–23*), they often focus on a narrow set of cell types and report conflicting results, suggesting the response is highly cell type dependent. There is a lack of quantitative data on how MNPs influence cell migration dynamics on different cell types. Moreover, many studies do not benchmark MNP exposure levels against concentrations detected in human tissues.

Investigations into the uptake of MNPs were mostly conducted using in vitro models under static conditions that fail to replicate the fluid properties of the physiological environment. In vivo, tissue cells are exposed to varying fluid viscosity, osmolarity, and hydrostatic pressure, which differ substantially from those in standard cell culture media (*24–27*). Among these factors, fluid viscosity has recently emerged as a critical regulator of particle uptake. Physiological fluid viscosity ranges from 1.5 to 8 centipoise (cP), notably higher than typical culture media (0.8 cP), and has been shown to modulate endocytosis and uptake of extracellular particles (*28, 29*). This regulation is mediated through mechanochemical crosstalk involving cytoskeletal elements such as actomyosin and ERM proteins, together with ion transporters like the Na^+^/H^+^ exchanger NHE1 (*28–33*). These cellular components sense and respond to extracellular fluid properties, generating water flux and cytoskeleton remodeling that ultimately alter cell surface mechanics and particle engulfment. Despite these insights, how such fluid-cell interactions influence MP uptake, retention, and release remains largely unexplored.

In this work, we investigate NP uptake in mice and in diverse cell types under physiologically relevant conditions, using quantitative microscopy and microfluidic transport assays to quantify cellular NP association, and examine how NP contamination affects cell proliferation and motility. The cell types studied span epithelial, endothelial, and fibroblast cells, as well as immune cells including primary human T cells. We find that NP contamination universally impairs cell proliferation, and produces cell type specific effects on cell motility. Immune cells, including healthy donor-derived CD4^+^ T cells, are particularly sensitive to NPs. Using a panel of pathway-specific inhibitors and transwell-based transport assays, we show that NP association, cellular retention, and transepithelial transport are regulated not only by endocytosis and exocytosis but also by ion transporters. Consistent with this, NP uptake is highly sensitive to extracellular fluid properties, particularly viscosity in the physiological range (1.5-8 cP). Finally, we reveal distinct NP types (e.g. PS, PE, and PP) interact with tissue cells. Together, our findings reveal the complex interactions between NPs and tissue cells and alterations in cell physiology.

## Results

### Quantifying cell-associated NPs and NPs retention in vitro and in vivo

To investigate how NPs associate with and are internalized by cells, we cultured cells with fluorescent carboxylated-polystyrene nanoplastic beads (0.1 µm in diameter, hereafter referred to as PS NPs unless otherwise noted) at varying concentrations and incubation times. This particle size permits association via both micro- and macropinocytosis pathways (*13*). Using membrane labeled Madin-Darby Canine Kidney II (MDCK II) epithelial monolayers, we visualized NP localization using confocal microscopy and *z*-stack imaging (Fig. 1A). After 6 hours of NP exposure followed by extensive PBS washes (*>*6 times), fluorescent signal was observed from both inside the cells (38.0%) and on the apical cell surface (62.0%). The surface-associated NPs appeared to strongly adhere to the plasma membrane. NP signals could be observed in cells undergoing mitosis, while no significant signals were observed in the nucleus or the cell-cell junctions. For clarity, we refer to the total of surface-adhered and internalized NPs as “cell-associated NPs”.

**Figure 1:**
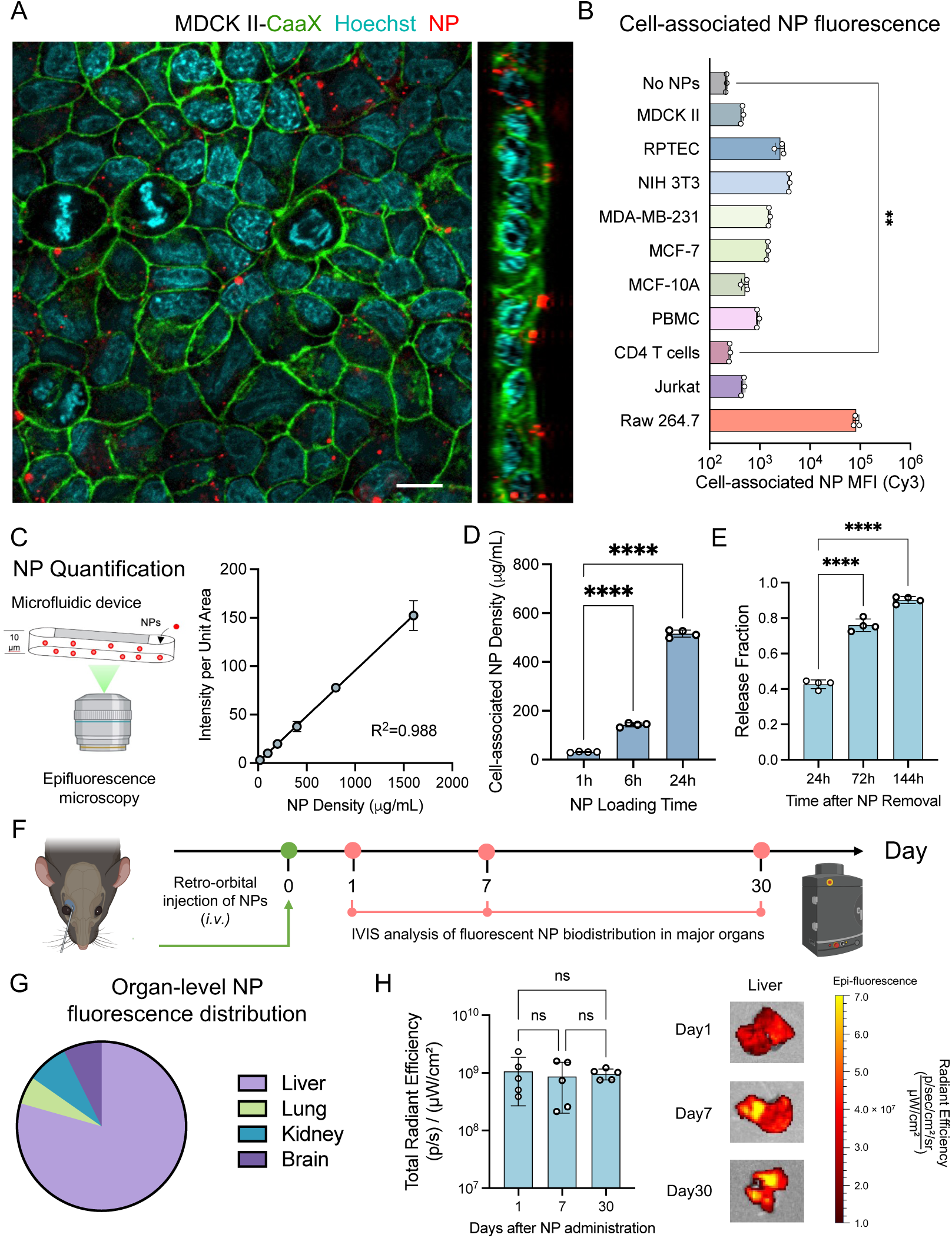
Quantification of Nanoplastics (NPs) association and release. (A) Representative confocal images of MDCK II monolayers expressing CaaX-GFP to label cell membrane (green), stained with Hoechst 33342 to label cell nucleus (cyan), and pre-incubated with 20 µg/mL red fluorescent NPs (red) for 24 hours. Panel on the right is a *z*-stack image. (B) Cell-associated NP fluorescence of different cell lines quantified using flow cytometry. The median fluorescence intensity (MFI) of N = 3 biological replicates is plotted in the log scale. (C) A method to quantify the mass of NPs using epifluorescence microscopy and microfluidic devices. 20 to 1600 µg/mL of NPs were injected into a microfluidic channel with known height similar to tissue cells, and imaged using an epifluorescence microscope. The total fluorescence per unit area is a linear function of the NP density. n = 3 technical repeats per condition. (D) Quantification of cell-associated NP density at different NP loading times in the MDCK II monolayer. 20 µg/mL NPs were loaded for 1, 6, and 24 hours, followed by washing and epifluorescence imaging. The mass density was then calculated based on the cell height measured in (A) and the calibration curve in (C). N = 4. (E) The relative NP release at different times after 24 hours of NP loading in the MDCK II monolayer. The release fraction is the loss of fluorescence intensity compared to the intensity measured immediately after NP loading by tracking the same region of cells. N = 4. (F) Schematic of the in vivo model. C57BL/6 mice were administered 100 µL of NPs (2 mg/mL) via intravenous (i.v.) injection, then sacrificed at 1, 7, and 30 days post-injection. NP retention was assessed by IVIS imaging. (G) NP distribution in major organs and brain 30 days post-injection. (H) Representative images and total radiant efficiency of NPs in liver at 1, 7, and 30 days post-injection. (B, C, D, E, H) Error bars represent standard deviation. Student’s *t*-test or one-way ANOVA was used. Representative statistical significance is shown: ns for *p >* 0.05, \**p <* 0.05, \*\**p <* 0.01, \*\*\**p <* 0.001, \*\*\*\**p <* 0.0001. (A) Scale bar = 10 µm.

We next measured NP association across a panel of cell types using flow cytometry (Fig. 1B and Fig. S1A). These included: two epithelial cell lines (MDCK and RPTEC/TERT1, an hTERT-immortalized human renal proximal tubule epithelial cells), a fibroblast cell line (NIH 3T3), three breast cancer cell lines with varying metastatic potential (MDA-MB-231, MCF-7, and MCF-10A), and immune cells including primary peripheral blood mononuclear cells (PBMCs), CD4 T cells, Jurkat T cells, and RAW 264.7 macrophages. All tested cell types showed significantly elevated NP signal compared to NP-free controls, indicating that NP uptake and association are universal among different cell types. CD4 T cells and Jurkat T cells exhibited the lowest median fluorescence intensity (MFI), while RAW 264.7 macrophages showed MFI levels 10-100 times higher than other cell types. Among the breast cancer lines, normal-like MCF-10A had lower NP association than both cancerous counterparts MCF-7 and MDA-MB-231.

To confirm that our experimental conditions corresponded to physiologically relevant NP levels, we quantified the absolute amount of cell-associated NPs using a precision microfluidic device (Fig. 1C). These microfluidic channels were designed with a height comparable to cell height (∼10 µm), which we measured accurately using confocal microscopy with point spread function deconvolution (*34*). At this scale, epifluorescence microscopy can capture all NP signal within the volume, enabling a calibration curve between NP density and fluorescence intensity (Fig. 1C). By imaging MDCK monolayers under the same conditions, we were able to calculate the absolute amount of cell-associated NPs. After 6 hours of loading at 20 µg/mL NP concentration, 142.6 µg/mL NPs were associated with MDCK cells. Given that mammalian cytoplasmic mass density is tightly regulated at ∼200 mg/mL (*34,35*), the estimated NP dry mass fraction is ∼0.07%. This fraction scales with loading concentration and duration (Fig. 1D and Fig. S1B). Reported NP levels in human tissues range from 60-5000 µg/g wet mass, which corresponds to an estimated 0.06-5% dry mass fraction based on a typical 1:10 dry-to-wet mass ratio. We thus conclude that our in vitro NP levels are within the physiologically relevant range.

We further investigated the retention and clearance of NPs from cells by tracking MDCK monolayers over time (Fig. 1E). Cells were loaded with NPs for 24 hours, washed thoroughly, and then incubated in NP-free media for up to 6 days. We imaged the same regions immediately after washing and at each subsequent time point, with daily media changes to prevent reattachment of extracellular NPs. Fluorescence intensity decreased by 42.7% (release fraction) in the first 24 hours post-wash, but 9.7% of the original signal remained even after 6 days (release fraction = 90.3%). These results suggest that NPs are not easily cleared from tissue cells, highlighting the potential for long-term intracellular retention.

Retention in vivo is a more complex scenario, as internalized NPs must travel in the circulatory system before being cleared. To determine whether NPs can persist within the body, we employed a mouse model. Fluorescent NPs were administered via intravenous (i.v.) injection at a total dose of 0.2 mg (2 mg/mL in 100 µL PBS), following previous protocols (*10, 11*)(Fig. 1F). At 1, 7, and 30 days post-injection, mice were sacrificed and NP distribution in major organs was quantified using an In Vivo Imaging System (IVIS). We observed fluorescent signals in liver, kidney, lung, and brain 30 days after NP administration. NPs were predominantly enriched in the liver (79.4%), one order of magnitude higher compared with kidney (7.91%), brain (7.37%), and lung (5.33%) (Fig. 1G). In contrast to the 90% NP release observed from monolayers in vitro within one week, NP retention in liver persisted within 30 days without statistically significant reduction (Fig. 1H). Together, these findings suggest slow NP clearance by tissue cells and persistent retention in vivo.

### NPs affect cell proliferation

Next, we examined how NP exposure influences cell proliferation. We found that growing cells with a high concentration of NPs (200 µg/mL) significantly reduced proliferation across all 9 cell types tested (Fig. 2A). At 20 µg/mL or lower, most cells proliferated at rates similar to the control. Immune cells, including primary CD4^+^ T cells, Jurkat T cells, and RAW 264.7 macrophages, appeared more sensitive to NP exposure, exhibiting significant reductions in proliferation even at 20 µg/mL. Long term exposure (*>* 48 hours) to NPs also significantly altered RAW 264.7 cell morphology (Fig. 2B). RAW 264.7 and NIH 3T3 proliferation results were validated by tracking the proliferation kinetics (Fig. 2C and D).

**Figure 2:**
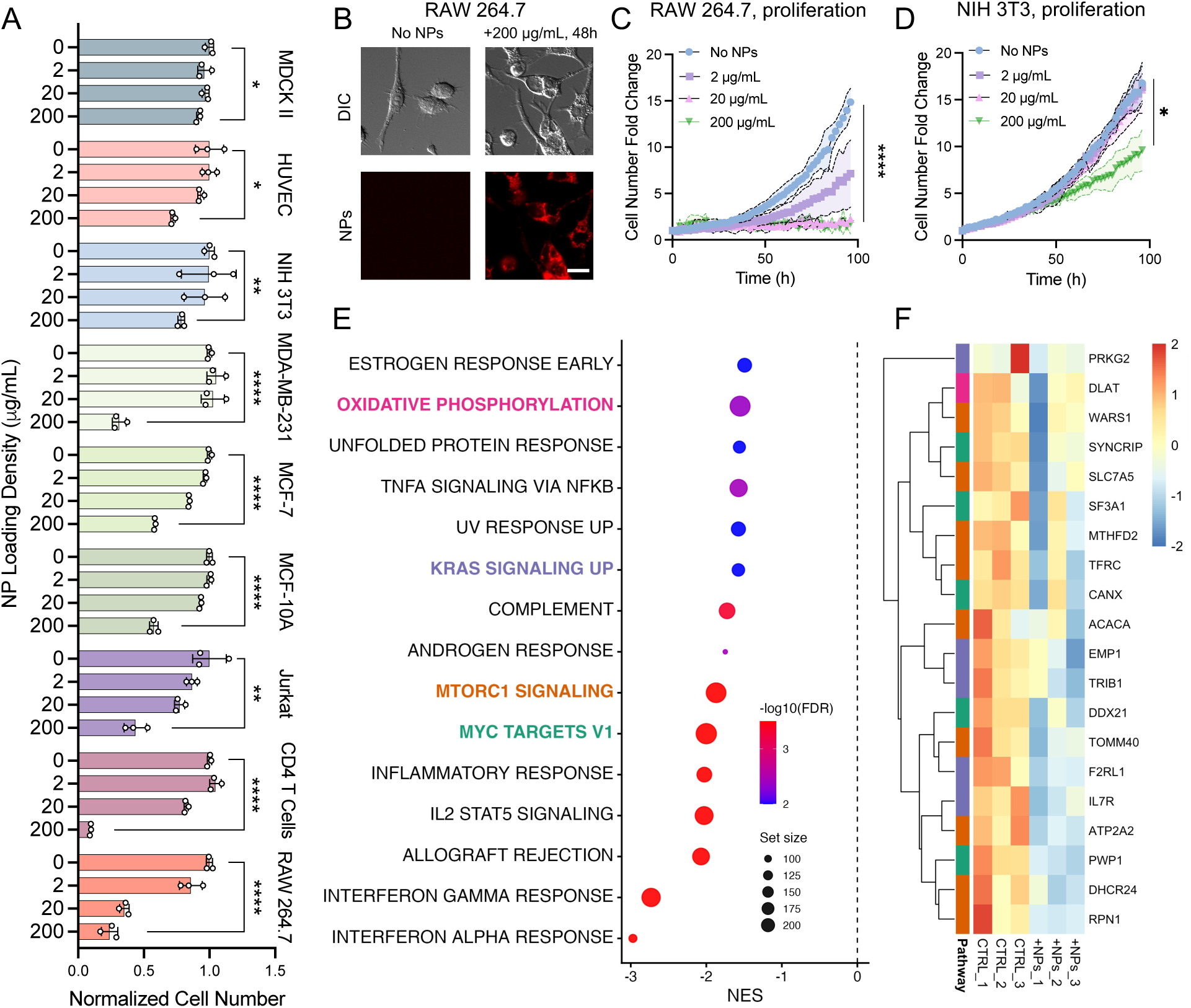
NPs affect cell growth. (A) Relative cell number fold changes of different cell types cultured with 0, 2, 20, or 200 µg/mL NPs for 4 days. Cell number was quantified by DNA staining using Hoechst 33342. N = 3. (B) Representative images of RAW 264.7 macrophages with and without NPs for 4 days. Red fluorescence indicates NPs. (C and D) Proliferation kinetics of RAW 264.7 macrophages (C) and NIH 3T3 fibroblasts (D) over 4 days, measured by cell number fold change using a quantitative phase microscope (Holomonitor) under varying NP concentrations as shown in (A). N = 3. (E) Gene set enrichment analysis (GSEA) of Hallmark pathways for NPs treated (+NPs; 50µg/mL for 48 hours) versus untreated control (CTRL) in CD4 T cells. X-axis denotes normalized enrichment score (NES). Points denote significantly enriched Hallmark gene sets, where point size reflects statistical significance (FDR-adjusted q value) and color reflects gene-set size. (F) Leading-edge expression heatmap for 4 selected Hallmark pathways enriched in +NPs versus CTRL colored in (E). Rows show leading-edge genes (core genes driving enrichment) and columns show biological replicates for CTRL and +NPs. Values are VST-normalized expression scaled per gene (row z-score). (A, C, and D) Error bars represent standard deviation. Student’s *t*-test was used. (C and D) Student’s *t*-test was conducted on the data at the last time point between untreated control and 200 µg/mL NPs. (B) Scale bar = 20 µm.

The observed proliferation reduction is in agreement with RNA sequencing results from primary CD4 T cells cultured with or without NPs for 2 days (+NPs versus CTRL). Although only a small set of individual genes are differentially expressed (Fig. S1D and E), pathway-level analysis revealed a coherent suppression of growth-associated programs. Specifically, gene set enrichment analysis (GSEA) (*36*) of MSigDB Hallmark gene sets (*37*) identified significant negative enrichment for MYC targets, mTORC1 signaling, KRAS signaling, and oxidative phosphorylation (Fig. 2E). Negative enrichment for these targets suggests a transcriptional shift away from biosynthetic and bioenergetic states that support cell-cycle progression (*38*). We therefore visualized leading-edge genes (gene subsets that contribute most strongly to the enrichment signal) from these growth-related Hallmark pathways across biological replicates (Fig. 2F). Because the normalized enrichment scores were predominantly negative in the +NPs versus CTRL contrast, the leading-edge genes were biased toward higher expression in the control condition relative to NP exposure. The resulting leading-edge signature included genes linked to nutrient uptake and anabolic metabolism (e.g., SLC7A5, ACACA, DHCR24), mitochondrial function and central carbon metabolism (e.g., DLAT, TOMM40), and ribosome biogenesis and RNA processing (e.g., DDX21, SF3A1, PWP1, SYNCRIP) (*39*). Notably, the expression of SLC7A5, an L-amino acid transporter that supports mTORC1 activation, is reduced (*40*). Together, these coordinated transcriptional changes upon NP exposure are consistent with diminished biosynthetic capacity and provide molecular support for the observed proliferation reduction.

### NPs affect cell motility and wound healing

We next examined whether NP exposure also affects cell motility and wound healing. In immune cells, both primary CD4 T cells and Jurkat T cells showed reduced migration speed after NP exposure in 3D collagen matrices, with longer exposure further decreasing migration speed (Fig. 3A and B). Mean squared displacement (MSD) analysis further revealed reduced migration persistence in Jurkat T cells after prolonged NP exposure (Fig. 3E and F). We also examined RAW 264.7 macrophages migrating on 2D surfaces. RAW 264.7 cannot sustain NP exposure for a long time (Fig. 2A-C), we thus chose to vary NP concentrations and reduced the exposure time (6 hours). NP exposure reduced RAW 264.7 migration speed only at high NP concentration, while the persistence increased and peaked at intermediate NP levels (Fig. 3C and G). We next examined a mesenchymal model, NIH 3T3 fibroblasts, where both migration speed and MSD progressively decreased with increasing NP concentration (Fig. 3D and H). Furthermore, single-cell analysis of NIH 3T3 revealed an inverse relationship between NP association and migration speed, with cells containing higher NP levels migrating more slowly (Fig. S2A).

**Figure 3:**
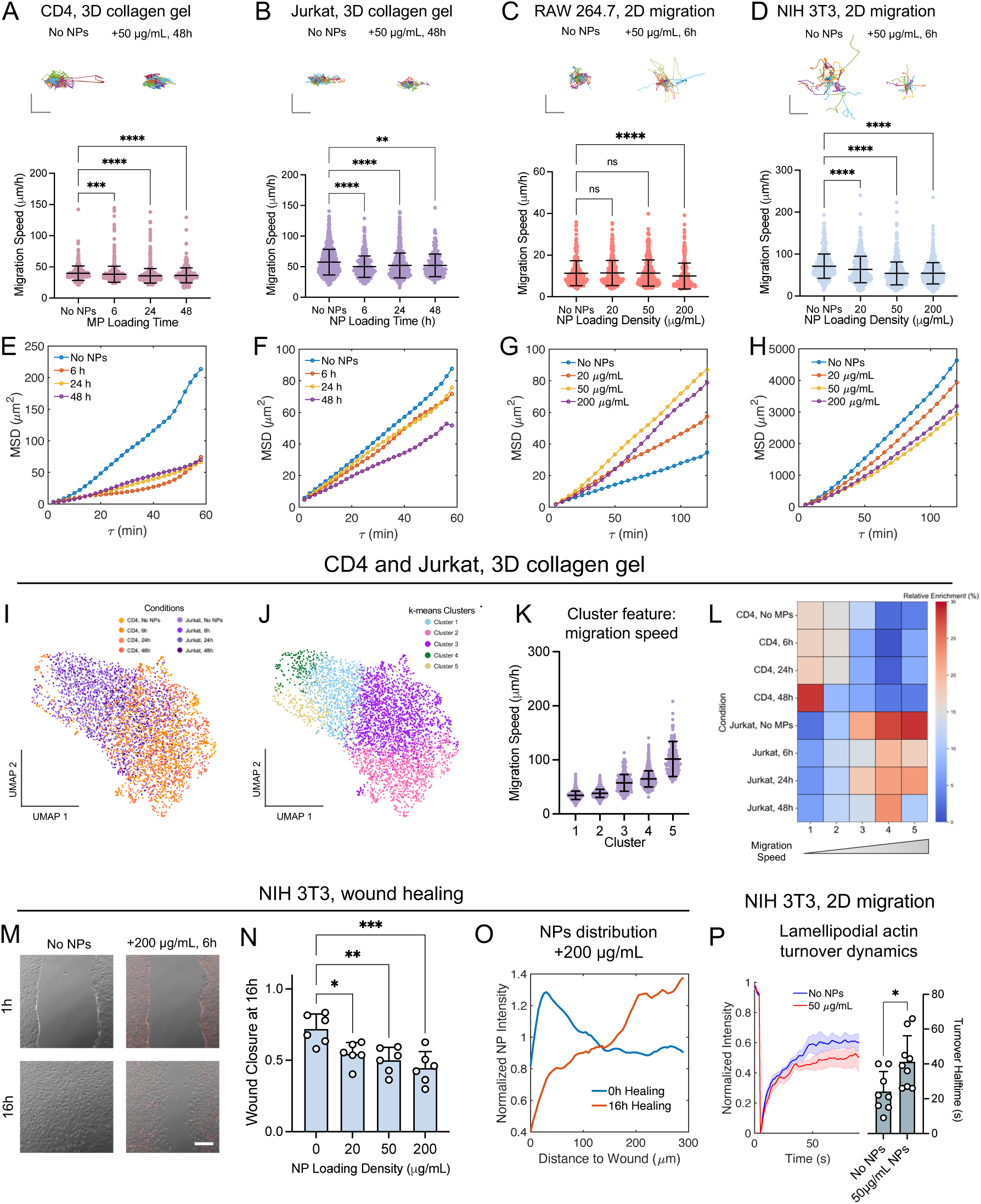
NPs affect cell motility and wound healing. (A and B) Representative migration trajectories and quantification of migration speed of CD4 T cells (A) and Jurkat T cells (B) in 3D collagen matrices. Cells were treated with 50 µg/mL NPs for 6, 24, or 48 hours, or left untreated. (C and D) Representative trajectories and migration speed of RAW 264.7 macrophages (C) and 3T3 fibroblasts (D) on collagen-coated 2D substrates. Cells were treated with 0, 20, 50, or 200 µg/mL NPs for 6 hours. RAW 264.7 were activated with Interleukin-6 (IL-6). (E-H) Mean squared displacement (MSD) of cells from panels A-D. (I) UMAP visualization of CD4 and Jurkat T cell trajectories under different NP exposure durations based on 79 motility parameters. (J) UMAP showing the same motility parameters, color-coded by clusters identified using unsupervised k-means clustering. (K) Average cell migration speed for each cluster shown in (J), ranked in ascending order. (L) Heatmap showing the relative enrichment of CD4 and Jurkat cells across different NP exposure durations within each cluster. Relative enrichment is defined as the proportion of cells from each condition within a given cluster. (M and N) Representative images (M) and quantification of relative wound closure at 16 hours (N) in NIH 3T3 fibroblast scratch-wound healing assays. N = 6. Cells were treated with 0, 20, 50, or 200 µg/mL NPs for 6 hours. NPs were removed before scratching and healing. (O) Distribution of cell-associated NP fluorescence in NIH 3T3 cells as a function of distance from the wound edge, measured immediately after scratch and 16 hours post-healing. (P) Actin turnover half-times in lamellipodia of EGFP–actin expressing NIH 3T3 cells with or without 50 µg/mL NPs for 6 hours1o1n collagen-coated 2D substrates. Turnover halftimes were quantified by Fluorescence Recovery After Photobleaching. n = 8 cells per condition. (A-D, N, and P) Error bars represent standard deviation. Shading represents the standard error of the mean. Kruskal-Wallis tests followed by Dunn’s multiple-comparison test were used. (A-D, and M) Scale bars = 15, 15, 5, 50, and 100 µm, respectively.

To further characterize NP-induced changes in cell migration behavior, we analyzed single-cell trajectories using the CaMI computational framework (*41, 42*). Unlike the average motility metrics shown in Fig. 3A-H, CaMI extracts multiple trajectory features to identify distinct migration phenotypic states through unsupervised clustering. In CD4 and Jurkat T cells migrating in 3D collagen matrices, CaMI identified five motility states shared across both cell types (Fig. 3I and J), wthere each state was characterized by distinct migration speeds (Fig. 3K). Regardless of their initial states, NP exposure shifted both cell types toward clusters with slower motility (Fig. 3L). A similar analysis of NIH 3T3 fibroblasts on 2D substrates identified four motility states, with NP exposure shifting cells toward lower motility clusters, consistent with the bulk migration results (Fig. S2F). In contrast, RAW 264.7 macrophages showed little change in the lowest or highest motility states but exhibited redistribution between intermediate clusters (Fig. S2E), consistent with their reduced migration speed but increased persistence.

We next extended our investigation from single-cell trajectories to collective migration by performing scratch wound healing assays in NIH 3T3 fibroblasts and epithelial monolayers (RPTEC and MDCK II). NP treatment reduced relative wound closure in NIH 3T3 fibroblasts, and cells migrating into the wound area exhibited visibly lower NP fluorescence (Fig. 3M and N). Spatial quantification of NP fluorescence also revealed a pronounced gradient, with markedly lower NP signal near the wound edge after healing (Fig. 3O). These findings are consistent with the single cell migration results, indicating that cells with lower NP association preferentially contribute to wound closure. In contrast, RPTEC and MDCK II epithelial monolayers showed no significant difference in wound closure dynamics, even at 200 µg/mL NP exposure and after extended incubation for up to 24 hours (Fig. S2B-D).

Interestingly, although NP exposure broadly altered cell motility, RNA sequencing of primary CD4 T cells did not reveal strong transcriptional changes in canonical migration-related pathways (Fig. 2E and F). This suggests that the reduced migration arises primarily from physical interactions between NPs and the cytoskeleton, the central driver of cell motility. In mesenchymal cells such as NIH 3T3 fibroblasts, actin filaments and associated regulatory proteins form highly dynamic networks at the cell leading edge (*14, 43*). We therefore hypothesized that NP exposure impairs migration by perturbing actin dynamics in this region. To test this, NIH 3T3 cells expressing EGFP–actin were analyzed using fluorescence recovery after photobleaching (FRAP) (*44–46*) (Fig. S2G). FRAP measurements in lamellipodia revealed an actin turnover half-time of ∼24.1 seconds in untreated cells (Fig. 3P), consistent with previous reports (*45*). Treatment with 50 µg/mL NPs for 6 hours increased the turnover half-time to ∼41.2 seconds, indicating slowed actin dynamics. These results support a model in which NP exposure mechanically interferes with actin network remodeling, thereby impairing cell migration.

Together, these results demonstrate that high-dose NP exposure universally impairs cell proliferation across diverse cell types, while effects on migration speed and persistence vary in a cell type-specific manner. These findings highlight the complex, context-dependent nature of NP-cell interactions and underscore the importance of considering both dosage and cell identity when evaluating the biological impact of MNP exposure.

### Endocytosis and ion transport regulate NP-cell association

To explore regulatory mechanisms governing NP-cell association, we used MDCK II epithelial monolayers and performed pharmacological perturbations to assess changes in NP uptake and release. Inhibition of clathrin-mediated endocytosis (dynasore) and myosin V–dependent intracellular trafficking (MyoVin-1) significantly reduced NP association in a dose-dependent manner (Fig. 4A and Fig. S3A,B), consistent with the known roles of endocytosis and intracellular trafficking in NP uptake (*13*). Inhibition of Rab GTPase-mediated trafficking pathways produced similar effects: CID-1067700 (Rab7 inhibitor), Nexinhib20 (Rab27 inhibitor), and fluvastatin (a statin that broadly disrupts Rab prenylation) all reduced NP association in MDCK II epithelial cells (Fig. 4A and Fig. S5A). Rab7 and Rab27 inhibition also reduced NP association in RPTEC epithelial cells and RAW 264.7 macrophages (Fig. S5B and D). In contrast, HUVEC endothelial cells showed distinct responses, with only a slight decrease after Rab7 inhibition and an unexpected increase following Rab27 inhibition (Fig. S5C).

**Figure 4:**
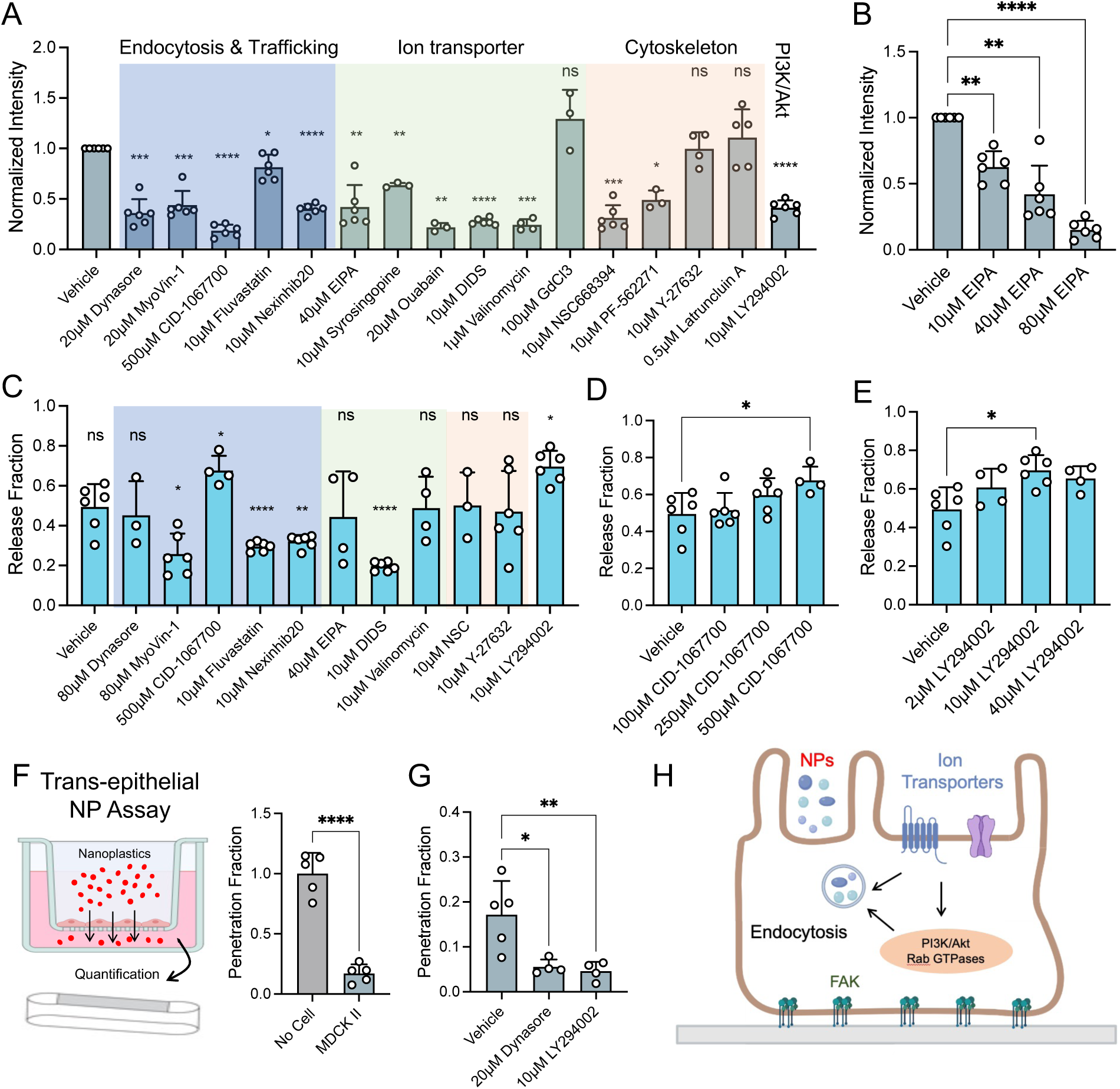
Endocytosis and ion transport affect NP-cell association. (A) Relative cell-associated NP fluorescence in MDCK II monolayers under treatment with different inhibitors. Cells were co-incubated with 20 µg/mL NPs and inhibitors for 6 hours. NP fluorescence intensity was normalized to vehicle control (DMSO). N = 3-6. (B) Dosedependent response of NP association under EIPA (NHE1 inhibitor) treatment. N = 6. (C) Relative NP release from MDCK II monolayers under various drug treatments. All samples were first incubated with 20 µg/mL NPs for 6 hours without drug treatment, followed by extensive washing and imaging to quantify the initial level of cell-associated NPs. Cells were then incubated in NP-free media containing inhibitors for 24 hours, after which NP fluorescence was measured again. The release fraction was calculated as the relative decrease in fluorescence intensity over this period. Conditions that compromised monolayer integrity due to cytotoxicity were excluded. N = 3-6. (D and E) Dose-dependent NP release fractions under treatment with CID-1067700 (Rab7 inhibitor; D) and LY294002 (PI3K inhibitor; E). N = 3-6. (F) Quantification of trans-epithelial NP transport sing a transwell system. MDCK II monolayers grown on 1 µm pore transwells were treated with 50 µg/mL NPs and inhibitors for 24 hours. NPs in the lower compartment were quantified using microfluidic devices. Empty transwells were used as baseline controls. N = 5. (G) Quantification of trans-epithelial NP transport in the presence of Dynasore or LY294002 using the assay in (F). N = 4-5. (H) Schematic illustration of pathways affecting NP-cell association. (A-G) Error bars represent standard deviation. (A and B) Paired *t*-test or RM one-way ANOVA was used. (C-G) Student’s *t*-test or one-way ANOVA was used.

Beyond classical endocytic regulators, endocytosis is closely coupled to dynamic remodeling of the cell surface. Recent work has shown that ion and water transport pathways, particularly those mediated by NHE1 and anion channels, regulate membrane swelling and boundary dynamics during processes such as endocytosis and cell shape change (*28, 34, 47–50*). These processes also involve the NHE1-binding partner ezrin and PI3K/Akt signaling cascades (*33, 51, 52*). Consistent with this mechanism, inhibiting NHE1 with EIPA, blocking anion transporters with DIDS, disrupting ezrin–actin binding with NSC668394, or inhibiting PI3K with LY294002 all reduced NP association in a dose-dependent manner (Fig. 4A, B and Fig. S3C,G,I). Perturbing additional ion transporters, including Na^+^/K^+^-ATPase (ouabain), monocarboxylate transporters MCT1/4 (syrosingopine), or increasing K^+^ permeability (valinomycin), produced a similar reduction (Fig. 4A and Fig. S3D,E). In contrast, disrupting important components of cell mechanics and mechanosensation by inhibiting stretch-activated Ca^2+^ channels with GdCl_3_, blocking ROCK with Y-27632, or depolymerizing actin with Latrunculin A had little effect, with the exception of focal adhesion kinase (FAK) inhibition using PF-562271 (Fig. S3F; Fig. 4A). Together, these results indicate that NP association is influenced by the interplay between membrane trafficking and ion transport, while the contribution of cell mechanics appears more complex.

We also investigated the regulatory role of these pathways in NP release. We first loaded cells with NPs in the absence of drugs, followed by a 24-hour release phase in the presence of each inhibitor. Surprisingly, most treatments that reduced initial NP association did not significantly alter NP release (Fig. 4C and Fig. S4). In particular, blocking clathrin-mediated endocytosis or ion transport pathways had little effect on the fraction of NPs released. Disrupting intracellular trafficking by inhibiting myosin V, Rab27, or statin decreased NP release (Fig. 4C; Fig. S4B; Fig. S6A). Two notable exceptions were observed with LY294002 and CID-1067700 (Fig. 4D,E), both of which reduced NP association but increased the released NPs (from 49.4% to 73.7% for LY294002 and to 67.6% for CID-1067700). While LY294002 had no significant effect on NP release in RPTEC, RAW 264.7, or HUVEC cells, CID-1067700 consistently increased NP release across all three cell types (Fig. S6B–D), indicating a broader role for Rab7-mediated trafficking in NP retention.

### NP transport across epithelial monolayers

In animal systems, NPs primarily enter the body via ingestion and must cross tissue barriers to reach internal organs. The epithelial monolayer, with tight cell-cell junctions (10 nm), poses a major barrier to NP transport. To assess whether NPs can traverse this barrier, we developed a transwell-based NP penetration assay. MDCK II cells were seeded into transwells with membranes with 1 µm pore size and cultured until confluence, verified by measuring trans-epithelial electrical resistance (TEER) (*53*). The small pore size prevents cell penetration across the membrane while allowing fluid and particle passage. NPs were added to the apical chamber for 24 hours, and the basolateral chamber was analyzed for NP content using our microfluidic quantification system. Compared to empty transwell controls, only 20% of NPs penetrated the MDCK II epithelial monolayer in 24 hours (Fig. 4F). No significant change in TEER was observed before and after NP exposure, suggesting that tight junction integrity remained intact (Fig. S1C). These findings suggest that while epithelial layers substantially restrict NP penetration, a fraction of NPs can still be transported across intact monolayers. By inhibiting Clathrin-mediated endocytosis or PI3K/Akt, the trans-epithelial NP transport was again effectively reduced (Fig. 4G). This supports the observation that NPs are actively transported through membrane endo- and exocytosis (Fig. 4H).

### Extracellular fluid physical properties influence NP association and release

Tissue cells in vivo experience fluid environments that differ significantly from standard in vitro culture conditions. Typical culture media have a low viscosity close to that of water (∼0.8 cP), while the viscosity of blood serum and interstitial fluid range from 1.5-8 cP (*24, 25, 28, 29*). To mimic these conditions, we increased media viscosity and examined NP association in MDCK II epithelial and HUVEC endothelial cells. We found that elevated fluid viscosity significantly enhanced NP association (Fig. 5 A-C). The maximum association occurred at 1.5 cP in MDCK II (∼ 4-fold increase) and at 1.5-2 cP in HUVEC (∼ 10-fold increase). Interestingly, NP association decreased beyond these peaks, revealing a non-monotonic response to viscosity. We further conducted a high-throughput flow cytometry screen of nine additional cell types (Fig. 5F). In all cases except one, elevated viscosity enhanced NP association, suggesting that viscosity is a broadly applicable regulator. The exception was RAW 264.7 macrophages, where increasing viscosity to 2 cP or 8 cP reduced NP association. This may be due to the already high baseline NP association in macrophages, which is 100-1000 times greater than in other cell types (Fig. 1B).

**Figure 5:**
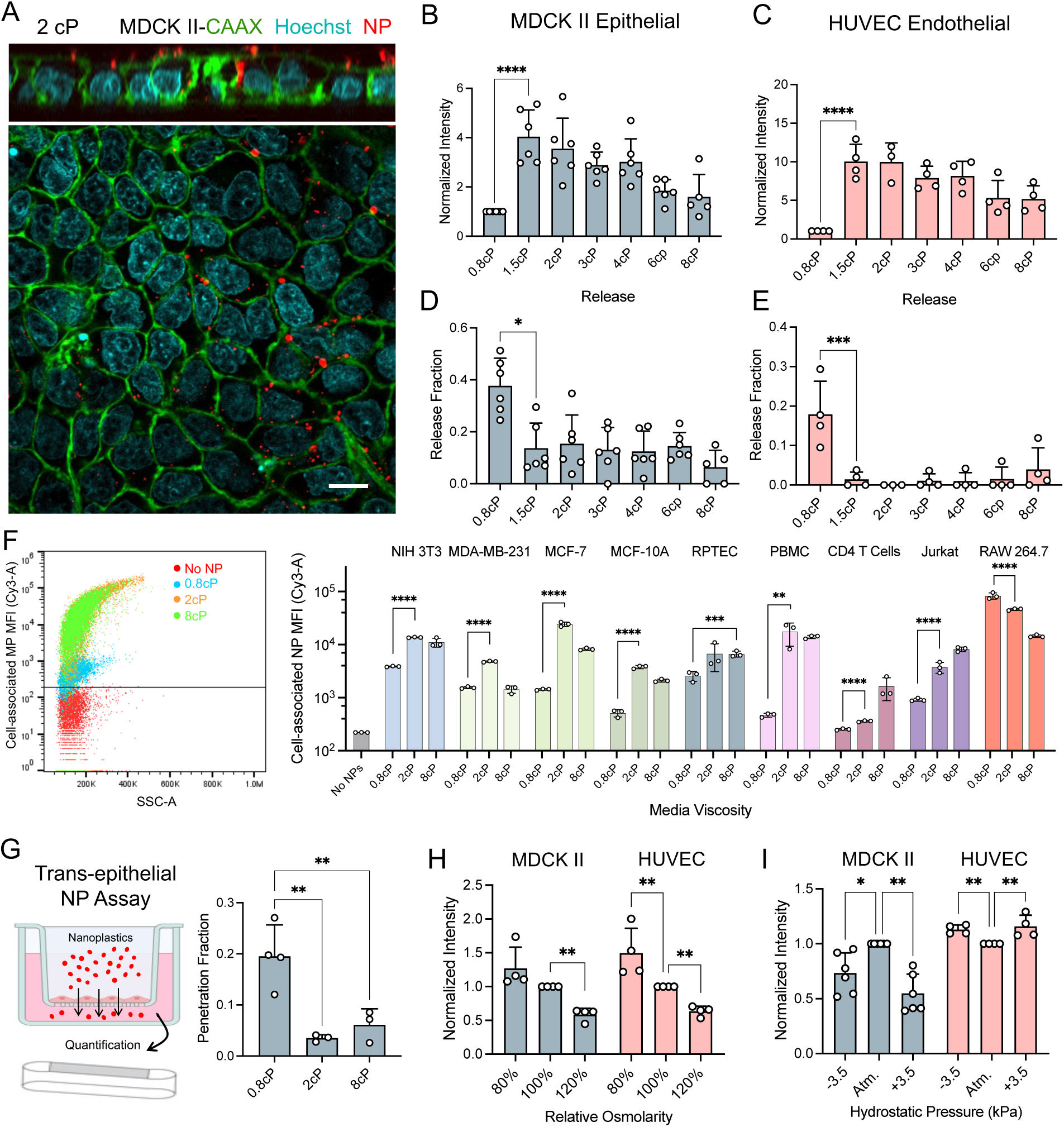
Fluid properties affect NP-cell association and release. (A) Representative confocal images of a MDCK II monolayer incubated with 20 µg/mL fluorescent NPs for 6 hours in 2 cP viscous media. Top panel shows a *z*-stack image. (B and C) Relative cell-associated NP fluorescence in MDCK II (B) and HUVEC (C) cells after 6-hour coincubation with 20 µg/mL NPs in media of varying viscosity. N = 3-6. (D and E) Relative NP release from MDCK II (D) and HUVEC (E) cells under different viscosity conditions. Cells were first loaded with 20 µg/mL NPs for 6 hours in 0.8 cP media, followed by washing and incubation in NP-free media of varying viscosity for 24 hours. NP fluorescence was measured before and after the release phase to calculate the release fraction. N = 3-6. (F) Left: a representative scatter plot of median fluorescence intensity (MFI) versus side scatter (SSC) of Jurkat cells treated with 20 µg/mL NPs for 6 hours at different viscosities. Right: MFI of cell-associated NPs in various cell lines treated with NPs in 0.8, 2, and 8 cP media. N = 3. (G) Quantification of trans-epithelial NP transport in MDCK II monolayers. Samples were treated with 50 µg/mL NPs in media of 0.8, 2, or 8 cP viscosity for 24 hours. The lower compartment contained media of the same viscosity. (H and I) Relative cell-associated NP fluorescence in MDCK II and HUVEC cells exposed to media of different osmolarity (H) or hydrostatic pressure (I). All samples were treated with 20 µg/mL NPs for 6 hours under the indicated fluid conditions. Pressure experiments were conducted in a sealed air-pressure chamber. N = 3-6. (B-I) Error bars represent standard deviation. Student’s *t*-test or one-way ANOVA was used. (A) Scale bar = 10 µm.

In addition to enhancing NP association, elevated fluid viscosity significantly impeded NP release. In both MDCK II and HUVEC monolayers, NP release after 24 hours was strongly suppressed under viscosities ranging from 1.5-8 cP (Fig. 5 D and E). In HUVECs, NP release was nearly undetectable under these conditions. Consistent with reduced NP release, we also observed a decrease in trans-epithelial NP transport when viscosity was elevated (Fig. 5 G). These observations suggest that the physiological level of fluid viscosity tends to trap NPs inside tissue cells and prevents trans-epithelial transport.

We next examined the roles of hydrostatic pressure and osmolarity in regulating NP association and release. Hydrostatic pressure varies between biological compartments, such as blood vessels, interstitium, and cerebrospinal fluid, in the range of a few kilopascals (*27*). Osmolarity is typically tightly regulated but can vary substantially in certain organs, such as the kidney. We found that exposing cells to a hydrostatic pressure gradient of as little as ±3.5 kPa significantly reduced NP association, without significantly altering the release fraction (Fig. 5I). On the contrary, both +/- 3.5kPa pressure rose NP association by ∼ 15% in HUVEC cells. Osmotic stress had opposing effects depending on direction. A 20% hypertonic shock increased NP association and decreased release, while a 20% hypotonic shock significantly reduced NP association and enhanced release (Fig. 5H).

Taken together, these results demonstrate that NP association and release are strongly influenced by the fluid microenvironment. Even modest deviations from standard in vitro conditions, such as changes in viscosity, pressure, or osmolarity, can result in substantial alterations in how tissue cells interact with NPs.

### Comparing association, release, and functional impacts of different NPs

So far, our experiments have focused on carboxylated-PS NPs, which are widely used in nanoplastic research due to their commercial availability and well-defined physicochemical properties (*10, 11,13*). Although PS NPs are prevalent in environmental samples, polyethylene (PE) and polypropylene (PP) MNPs are reported to be among the most abundant polymers detected in human tissues (*3–7*). Compared with negatively charged, relatively hydrophilic carboxylated-PS NPs, PE and PP NPs are largely neutral, more hydrophobic, and possess lower buoyant density (Fig. 6A). We therefore quantified the association and release of PE and PP NPs with the same size and shape as PS NPs in MDCK II cells. Similar to PS NPs, cell-associated PE and PP NPs increased with longer incubation time, although their absolute cellular association levels were lower than those observed for PS (Fig. 6B and Fig. S7A and B). In contrast, a larger fraction of PE and PP NPs was released within 24 hours after removal of extracellular particles compared with PS NPs (Fig. 6C). The association and release dynamics were similar between PE and PP.

**Figure 6:**
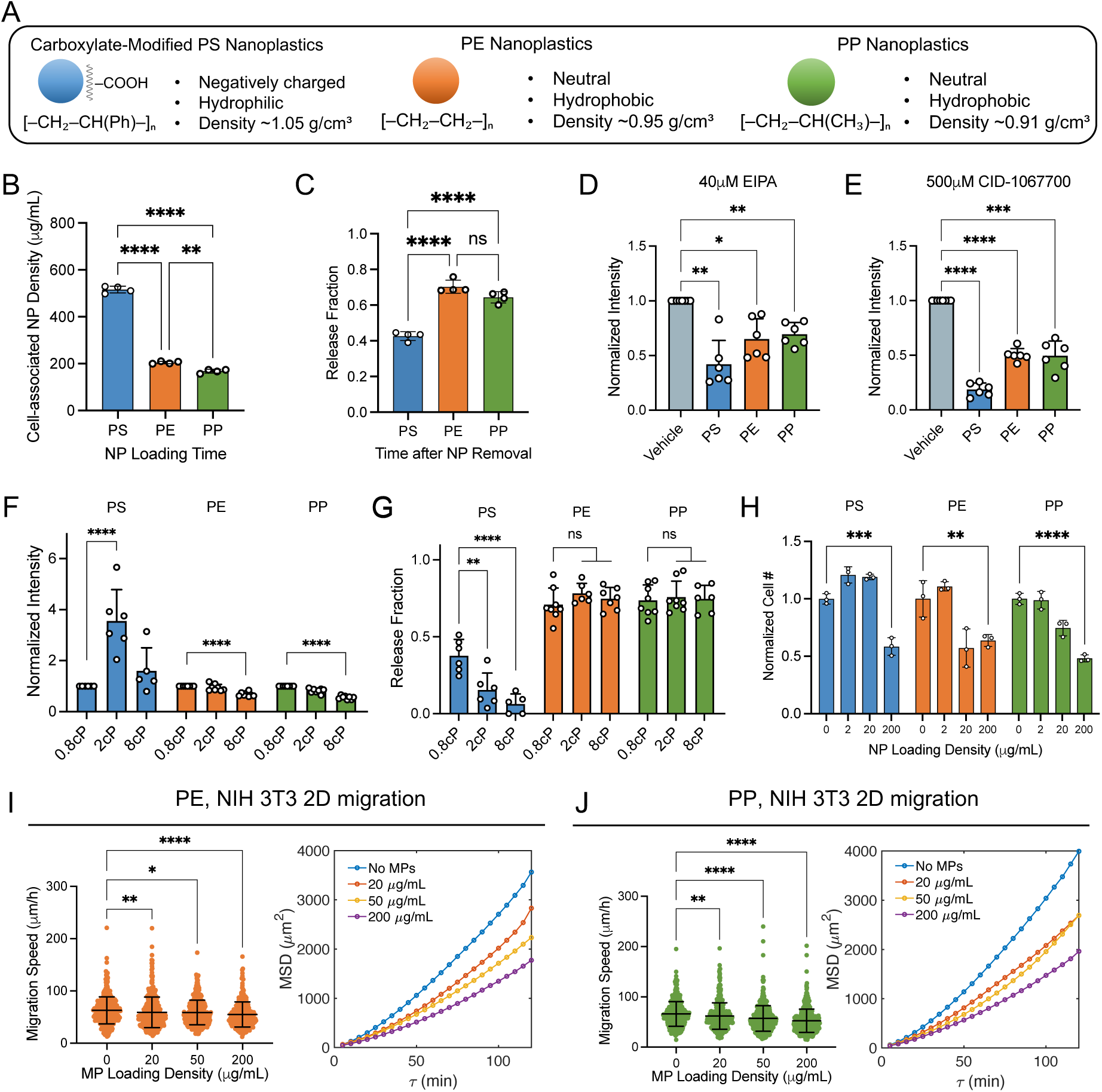
Comparing the association, release, and functional impacts of carboxylated-PS, PE, and PP NPs. (A) Schematic overview of carboxylate-modified PS, PE, and PP NPs. (B and C) Quantification of cell-associated PS, PE, and PP NP density (B) after loading MDCK II monolayers with 20 µg/mL NPs for 24 hours, and NP release 24 hours after NP removal (C), following the method described in Fig. 1D and E. N = 4. (D and E) PS, PE, and PP NP association in MDCK II monolayers treated with EIPA (NHE1 inhibitor; D) or CID-1067700 (Rab7 inhibitor; E). N = 6. (F and G) NP association (F) and release fraction (G) under different viscosity conditions in MDCK II monolayers. (H) Relative cell number fold changes of NIH 3T3 cells cultured with 0, 2, 20, or 200 µg/mL NPs for 4 days. Cell number was quantified by DNA staining using Hoechst 33342. N = 3. (I and J) Migration speed and mean square displacement (MSD) of 3T3 fibroblasts on collagen-coated 2D substrates treated with 0, 20, 50, or 200 µg/mL PE (I) and PP (J) NPs for 6 hours. N = 3. (B-J) Error bars represent standard deviation. One-way ANOVA or Student’s *t*-test was used. RM one-way ANOVA was used in (D-F).

We next examined whether these different NPs share similar association pathways. Inhibition of NHE1 using EIPA, clathrin-mediated endocytosis using dynasore, and Rab7-mediated intracellular trafficking using CID-1067700 reduced NP association across all plastic types tested (Fig. 6D, E and Fig. S7C). However, inhibition of PI3K/Akt signaling using LY294002 did not significantly affect PE or PP NP association (Fig. S7 D). Moreover, while CID-1067700 and LY294002 increased the release of PS NPs, neither treatment altered the release of PE or PP NPs (Fig. S7E and F). In addition, while association and release of PS NPs are highly sensitive to extracellular fluid viscosity, it had minimal effect on PE and PP NPs (Fig. 6F and G). At 8 cP, we even observed a slight reduction in PE and PP NP association. Together, these results indicate that NP–cell interactions are polymer-type dependent.

Finally, we asked whether different NPs have distinct functional impacts on cells. NIH 3T3 fibroblasts were highly sensitive to NP exposure, showing a significant reduction in proliferation and decreased motility. Therefore, we used NIH 3T3 cells as the model cell line to compare the functional effects of different NP types. At a moderate dose (20 µg/mL), PS NPs did not significantly affect NIH 3T3 proliferation, whereas PE and PP NPs at the same concentration had already reduced cell number after 4 days (Fig. 6H). At a higher dose (200 µg/mL), all NP types caused similar reductions in proliferation. In single-cell migration assays, PE and PP NPs produced effects similar to PS NPs, with both migration speed and persistence decreasing as NP dose increased (Fig. 6I and J). Overall, PE and PP NPs have similar effects to PS NPs on cell proliferation and motility.

## Discussion

In this work, we systematically investigated how NPs affect cell proliferation and motility across a range of cell types commonly used in biomedical studies. To ensure physiological relevance, we developed a microfluidics-based assay combined with fluorescence microscopy to quantify the concentration of cell-associated NPs. While NP exposure consistently reduced cell proliferation at high doses, its effects on motility were more nuanced and highly cell type dependent. We further identified that NP-cell association is regulated not only by endocytosis but also by ion transporters and the extracellular fluid environment, particularly fluid viscosity. The significant changes in NP-cell association observed under physiologically relevant fluid viscosities raise important considerations about how in vitro assays are designed and interpreted under standard low-viscosity culture conditions.

Our mouse model further suggests that NPs introduced into the systemic circulation can persist in vivo for extended periods. While our in vitro assays demonstrate mechanisms of NP-cell association and reveal routes for NP release from tissue cells, strategies to enhance systemic NP clearance in vivo remain largely unexplored. Systemic clearance in vivo likely requires coordinated transport through the circulatory and reticuloendothelial systems. Notably, despite the strong effects of Rab7 inhibition (CID-1067700) on NP association and release in vitro, preliminary mouse studies using both preventive and post-exposure treatment paradigms showed limited impact on overall NP retention (Fig. S8), highlighting the complexity of in vivo clearance mechanisms.

A broader implication of these findings is that MNP behavior may depend strongly on MNP identity. The relative composition of microplastics found in human tissues (*3, 4, 7*) does not reflect their environmental abundance (*2, 54*), suggesting selective accumulation or differential transport mechanisms among MNP types (*17*). This could arise from differences in size, shape, surface charge, or other physical properties. In this study, we used commercially available microbeads with well-defined spherical shape and 100 nm diameter. While useful for controlled experiments, these particles do not fully represent the diversity of micro- and nanoplastics encountered in natural or clinical settings. Although the influence of microplastic size is relatively well studied, the effects of shape and surface charge have only recently gained attention (*55*). Many studies use commercially available PS beads, often with surface carboxylation to improve colloidal stability and minimize aggregation. However, this modification substantially alters surface charge and interfacial properties. Our comparison indicates that carboxylated PS beads exhibit context-dependent differences in cell interactions relative to similarly sized PE and PP particles, while producing broadly similar effects on proliferation and motility. These findings underscore the need to systematically investigate the biological impact of different MNP polymer identities.

Our data also raise mechanistic questions about how micro- and nanoplastics associate with cells beyond classical endocytic pathways. We identified roles for several ion transporters and the PI3K/Akt pathway in modulating MNP association, but their exact contributions remain unclear. For example, do these regulators directly influence endocytosis, or do they act indirectly through changes in membrane tension, cytoskeletal organization, or cellular energy metabolism? Na^+^/K^+^- ATPase and PI3K/Akt are both known to affect cellular bioenergetics and signaling, raising the possibility that reduced MNP association under their inhibition may arise from broader changes in metabolic state.

## Materials and Methods

### Cell Culture

NIH 3T3, Raw264.7, and Jurkat cells were purchased from the American Type Culture Collection (ATCC). MDCK and HUVEC cells were a gift from Denis Wirtz (Johns Hopkins University, Baltimore, MD). MCF-10A, MCF-7, and MDA-MB-231 cells were a gift from Konstantinos Konstantopoulos (Johns Hopkins University, Baltimore, MD). NIH 3T3, MDCK, MCF-7, MDA-MB-231, and Raw264.7 cells were cultured in Dulbecco’s Modified Eagle Medium (DMEM; Corning) supplemented with 10% fetal bovine serum (FBS; Sigma) and 1% antibiotic solution containing 10,000 units/mL penicillin and 10,000 *µ*g/mL streptomycin (p/s; Gibco). MCF-10A cells were cultured in DMEM/F-12 supplemented with 5% horse serum, 20 ng/mL epidermal growth factor, 0.5 *µ*g/mL hydrocortisone, 100 ng/mL cholera toxin, 10 *µ*g/mL insulin, and 1% p/s, as previously described (*56*). HUVEC cells were cultured in Vascular Cell Basal Medium supplemented with Endothelial Cell Growth Kit - VEGF (ATCC). Jurkat cells were cultured in RPMI 1640 (Gibco) supplemented with 10% FBS and 1% p/s. All adherent cells were passaged using 0.05% trypsin-EDTA (Gibco).

Human peripheral blood mononuclear cells (PBMCs) were isolated from peripheral blood obtained from a single de-identified leukopak donor using density-dependent centrifugation. Leukopak blood was obtained from the Johns Hopkins Center for Genomic Medicine. CD4^+^ T cells were purified from the PBMC population using a magnetic bead-based negative isolation kit (STEMCELL Technologies). Both PBMCs and CD4^+^ T cells were resuspended in complete RPMI medium containing 10% FBS, 1% sodium pyruvate, 1% HEPES, 1% Pen/Strep (all from Gibco), and 4 ng/mL beta-mercaptoethanol (Sigma-Aldrich). To activate CD4^+^ T cells, isolated cells were cultured at a density of 1×10^6^ cells/mL in medium containing 25 *µ*L/mL anti-CD3/CD28 Dynabeads (Thermo) and 50 ng/mL human recombinant IL-2 (STEMCELL Technologies) for 7 days, followed by 7 days without beads to allow for rest and expansion.

All cell cultures and live-cell experiments were conducted at 37^◦^C in a humidified atmosphere with 5% CO_2_.

### Nanoplastics and Pharmacological Inhibitors

Carboxylated polystyrene (FluoSphere, Invitrogen), polyethylene (Biotyscience Inc), and polypropylene (Biotyscience Inc) microbeads with a diameter of 0.1 µm and fluorescence excitation/emission at 580 nm/605 nm were used as nanoplastics (NPs) in this study. NP beads were vortexed thoroughly before each use. In selected experiments, cells were treated with the following pharmacological agents: vehicle control (0.25% DMSO; Invitrogen), Dynasore (Sigma), MyoVin-1 (Sigma), CID-1067700 (MilliporeSigma), EIPA (Tocris), syrosingopine (Cayman Chemical), Ouabain (Tocris), Valinomycin (Tocris), Gadolinium chloride (GdCl_3_; Sigma), NSC668394 (MilliporeSigma), PF-562271 (Selleckchem), Y-27632 (Tocris), and LY-294002 hydrochloride (Sigma). To investigate regulatory pathways involved in NP association, inhibitors were co-administered with NPs for 6 hours. To investigate pathways involved in NP release, cells were first incubated with NPs alone for 6 hours, followed by washing and subsequent treatment with inhibitors for 24 hours.

### Epi-fluorescence and Confocal Microscopy

For epi-fluorescence and phase-contrast imaging, a Zeiss Axio Observer inverted wide-field microscope was used, equipped with either a 20× air, 0.8-NA objective (for glass-bottom dishes) or a 20× air, 0.4-NA long-working-distance objective (for plastic dishes), and an Axiocam 560 mono CCD camera. In some experiments, a similar setup equipped with a Hamamatsu Flash4.0 V3 sCMOS camera was used.

For confocal imaging, a Zeiss LSM 800 confocal microscope equipped with a 63× oil-immersion, 1.2-NA objective was used. All microscopes were outfitted with a CO_2_ Module S (Zeiss) and a TempModule S stage-top incubator (Pecon), set to 37 ^◦^C and 5% CO_2_ for live-cell imaging. ZEN 2.6 or ZEN 3.6 software (Zeiss) was used for image acquisition.

### Live Cell Reporters, Lentivirus Preparation, and Transduction

MDCK and NIH 3T3 cells stably expressing pHR GFP-CaaX (Addgene #113020) were generated via lentiviral transduction and selected by flow cytometry. For lentivirus production, HEK 293T/17 cells were co-transfected with psPAX2, VSVG, and the lentiviral plasmid of interest. Forty-eight hours after transfection, viral supernatant was harvested and concentrated by centrifugation. Wildtype MDCK or NIH 3T3 cells at 60-80% confluency were incubated for 24 hours with 100× concentrated virus suspension and 8 *µ*g/mL Polybrene Transfection Reagent (MilliporeSigma). NIH 3T3 cells expressing EGFP-actin (Addgene #56421) were generated using Lipofectamine 3000 (Invitrogen) following manufacturer’s protocol.

### Confocal Imaging of NPs in MDCK Cells

50,000 MDCK-GFP-CaaX cells per well were seeded into glass-bottom 24-well plates (Cellvis) pre-coated with 50 µg/mL Collagen-I (Enzo) and incubated at 37 ^◦^C for 1 hour. Cells were grown to confluence, followed by incubation with 20 µg/mL NPs for 24 hours in either normal media or 2 cP media. Prior to imaging, samples were washed at least six times with DPBS to remove all non-associated NPs. Cells were then stained with Hoechst 33342 (1:2000 dilution) for 15 minutes at 37 ^◦^C, followed by an additional three washes with DPBS. Samples were imaged using confocal microscopy with Z-stack acquisition.

### NPs Measurement Using Flow Cytometry

50,000 adherent cells or 200,000 suspension cells per well were seeded into 24-well plates and incubated for 24 hours prior to treatment. 20 µg/mL NPs were added as described above. Four hours after NP administration, cells were collected, washed twice with ice-cold PBS, and stained with LIVE/DEAD™ Fixable Aqua viability dye (Thermo Fisher Scientific, Cat# L34966) according to the manufacturer’s protocol. Fluorescence data were acquired using an Attune NxT flow cytometer (Thermo Fisher Scientific) and analyzed in FlowJo v10.8.1. The gating strategy used for analysis is detailed in Fig. S1. In selected experiments testing the effect of fluid viscosity, culture media of defined viscosities were prepared and maintained throughout the 4 four NP administration, as described in the corresponding section below.

### Quantification of NP Association and Release

Microfluidic channels with defined channel heights were fabricated as described below to generate a calibration curve relating NP density to fluorescence intensity. NPs were suspended in DMEM at various concentrations and injected into the channels, followed by imaging using the epi-fluorescence microscope described above. Control channels containing only DMEM were imaged to establish background signal. After subtracting background fluorescence, total intensity was summed over the entire image. The epi-fluorescence microscope captured signal across the full channel depth, as shown in our previous work (*33*), enabling correlation between NP mass (or mass density) and fluorescence intensity.

50,000 MDCK cells per well were seeded into glass-bottom 24-well plates (Cellvis) pre-coated with 50 µg/mL Collagen-I (Enzo) and incubated at 37 ^◦^C for 1 hour. Cells reached confluence and formed a monolayer within 2-3 days. NPs at varying concentrations were added to each well for different durations, as described in the main text. Prior to imaging, cells were washed three times with DPBS. Epi-fluorescence imaging and corresponding background images (DMEM without NPs) were acquired under identical conditions. After background subtraction, total image intensity was summed. Cells were then incubated in NP-free media for an additional 24 hours. After three more washes with DPBS, the same regions were re-imaged to quantify NP release. In parallel, the average height of the MDCK monolayer was measured using confocal microscopy and found to be approximately 9 µm, closely matching the microfluidic channel height. Using the previously obtained calibration curve, the absolute NP mass density was calculated.

For high-throughput NP screening, 50,000 MDCK or 30,000 HUVEC cells per dish were seeded into tissue culture-treated plastic dishes (Fisher). Cells reached confluence within 2-3 days. Imaging and data analysis followed the same procedures as described above, using a 20×, 0.4-NA long working distance objective. All experiments included a parallel control condition, and fold changes relative to the control were calculated as the normalized intensity for cell-associated NPs. Cells were then incubated in NP-free media for 24 hours and re-imaged in the same regions following three DPBS washes. Relative changes in fluorescence intensity were calculated to determine the release fraction.

### IVIS Imaging for Quantification of Fluorescence-Labeled NP Biodistribution in Mice

Ex vivo fluorescence imaging was performed to evaluate organ-specific biodistribution of fluorescently labeled NPs. At the designated time point post-administration, C57BL/6 mice were euthanized by CO inhalation in accordance with institutional animal care guidelines. Major organs, including the heart, liver, spleen, lungs, and kidneys, were harvested immediately and rinsed with PBS to remove residual blood. Organs were imaged using an IVIS Spectrum imaging system with appropriate excitation and emission filter settings for Cy3 fluorescence. Fluorescence intensity was quantified as radiant efficiency using Living Image software (Living Image^®^ 4.4). For experiments assessing the prophylactic and therapeutic effects of CID-1067700 on NP clearance, 16 mg/kg CID-1067700 dissolved in 50 µL DMSO was administered by intraperitoneal (i.p.) injection either 1 hour before NP administration (prophylactic assessment) or 1 day after NP administration (therapeutic assessment). Tolerability of this dose has been previously established (*57*). Mice were sacrificed 7 days after injection.

### Trans-epithelial NPs Assay and Quantification

50,000 MDCK cells per insert were seeded into collagen-coated 1.2 mm PET transwells with 1 µm pore size (Corning). Cells were allowed to grow for 2-3 days to reach confluence. Confluence was confirmed by measuring trans-epithelial electrical resistance (TEER) using an EVOM-2 system (WPI), following previously described protocols (*53*). 50 µg/mL NPs were added to the apical compartment of each transwell and incubated for 24 hours. Media from the basal compartment was collected, and NP concentration was quantified using the microfluidic-based fluorescence method described above. A control experiment without cells was performed using collagen-coated transwells to determine the baseline NP transport rate. Following the assay, TEER was measured again to confirm that epithelial confluence was maintained throughout the experiment.

### Microfluidic Device Fabrication

Microfluidic channel masks were designed using AutoCAD and fabricated by FineLineImaging. Silicon molds were produced using SU8-3010 photoresist (Kayaku) following standard photolithography procedures and the manufacturer’s protocol. Two photoresist layers were sequentially spin-coated onto silicon wafers (IWS): the first at 500 rpm for 7 seconds with an acceleration of 100 rpm/s, and the second at 2,000 rpm for 30 seconds with an acceleration of 300 rpm/s. A 4 minute soft bake was performed at 95 ^◦^C, followed by UV exposure to pattern the negative photoresist. Final channel dimensions were 16 mm in length and 1.2 mm in width. PDMS elastomer and curing agent (Sylgard 184, Dow Corning) were mixed at a 10:1 ratio, stirred vigorously, degassed under vacuum, poured over the silicon wafer, and cured at 80 ^◦^C for 45 minutes. Devices were cut to final dimensions using a razor blade, and inlet/outlet ports were punched with a blunt-tipped 21-gauge needle (McMaster-Carr). Devices were cleaned by sonicating in 100% isopropyl alcohol for 10 minutes and dried using compressed air. PDMS devices and sterilized glass-bottom 6-well plates (Cellvis) were exposed to oxygen plasma for 1 minute to enable bonding. Bonded devices were placed in an oven at 80 ^◦^C for 45 minutes to reinforce adhesion. Final channel height was approximately 9 µm, and exact heights were measured for each device using confocal microscopy.

### Cell proliferation assays

We used two independent methods to assess the effect of NPs on cell proliferation. In the first method, 5,000 cells were seeded into plastic 24-well plates (Falcon) 1 day prior to NP treatment. Cells were incubated for 4 days with 500 µL of media containing 0, 2, 20, or 200 µg/mL NPs. The commercial NP stock solution contained sodium azide as a preservative. The final concentration of sodium azide in the media is low, but may still affect cell proliferation at long term. To avoid this, the sodium azide concentration was adjusted to 20 µM across all conditions. After 4 days, NPs were removed, and samples were washed with DPBS before staining with Hoechst 33342 at 37 ^◦^C for 15 minutes (1:2000 in DPBS, Invitrogen). Total DNA content, used as a proxy for cell number, was quantified by measuring Hoechst fluorescence intensity using a plate reader (SpectraMax M3).

In selected experiments, cell proliferation kinetics were measured using a Holomonitor M4 quantitative phase imaging system (PHI). 5,000 cells were seeded into 24-well optical-bottom plates (lumox multiwell, SARSTEDT) 1 day prior to NP treatment. Cells were then treated with NPs following the same protocol as above. The Holomonitor captures phase shifts in transmitted light caused by differences in refractive index between cells and the surrounding medium. Phase images were acquired at regular intervals, and cell numbers were quantified using the Holomonitor M4 analysis software. Individual cells were segmented based on a height threshold and minimum spreading area, and proliferation was determined by calculating the average number of cells per frame over time.

### RNA sequencing and pathway enrichment analysis

Primary human CD4^+^ T cells were isolated and activated as mentioned above, and cultured without (CTRL) or with 50 µg/mL NPs (+NPs) for 2 days before RNA extraction for bulk RNA sequencing. Total RNA was extracted and purified using RNeasy Mini Kit (Qiagen) following the manufacturer’s protocol. Non-stranded mRNA-seq libraries were prepared using the VAHTS Universal V10 RNA-seq Library Prep Kit for Illumina (Vazyme). Poly(A)+ mRNA enrichment was performed using poly-T oligo–attached magnetic beads. Following fragmentation, first-strand cDNA synthesis was performed using random hexamer priming, followed by second-strand cDNA synthesis. Libraries were constructed through end repair, A-tailing, adapter ligation, size selection, PCR amplification, and purification according to the manufacturer’s protocol. Libraries were quantified using Qubit and real-time PCR and assessed for fragment size distribution using a Bioanalyzer. Pooled libraries were sequenced on an Illumina NovaSeq X Plus instrument using 150 bp paired-end reads (PE150).

Differential expression between +NPs and CTRL was performed with DESeq2 (*58*), using a negative binomial generalized linear model with condition as the primary factor. Gene-wise dispersion was estimated by DESeq2 and shrunken log_2_ fold-changes and Wald statistics were obtained for the +NPs versus CTRL contrast. Multiple testing correction was performed using the Benjamini–Hochberg procedure to control the false discovery rate (FDR), and genes were considered differentially expressed based on adjusted *p*-value and effect-size thresholds as indicated in the main text. To assess coordinated transcriptional changes beyond individual differentially expressed genes, gene set enrichment analysis (GSEA) (*36*) was performed using the MSigDB Hallmark gene sets (*37*). Genes were ranked genome-wide by the signed DESeq2 Wald statistic for the +NPs versus CTRL contrast, and enrichment was computed with a pre-ranked approach. Normalized enrichment scores (NES) and FDR-adjusted significance values were reported for each Hallmark gene set. For visualization and interpretability, leading-edge genes were extracted for selected growth-related Hallmark pathways. A compact leading-edge heatmap was generated from variance-stabilized transformed expression values (VST) from the DESeq2 object. For the heatmap, VST expression was scaled per gene (row-wise *z*-score) across samples, and genes were annotated by their corresponding Hallmark pathway membership.

### Scratch-Wound Healing Assay

50,000 3T3, MDCK, or RPTEC cells per well were seeded into collagen-coated 24-well plates as described above. Cells were grown for 2-3 days to reach confluence (4-5 days for RPTEC cells). Prior to wounding, 500 µL of media containing 0, 20, 50, or 200 µg/mL NPs was added to each well and incubated for 6 hours (for 3T3) or 24 hours (for MDCK and RPTEC), followed by three washes with PBS. Sodium azide concentration was adjusted to 20 µM across all conditions during NP administration as described above. Wounds were created by scratching the monolayer using a 200 µL pipette tip. Wound closure was monitored for 16 hours using phase-contrast microscopy. Images were analyzed manually in Fiji (ImageJ) to quantify the wound area, and relative wound closure speeds were calculated.

### Cell Migration Assay in 3D Collagen Gel and Data Analysis

To form hydrogels, rat tail type I collagen (Corning) was neutralized by mixing with 0.2 N NaOH to adjust the pH to 7-7.5. The neutralized solution was diluted to 0.5 mg/mL using a mixture of 10× HBSS (Sigma-Aldrich) and complete RPMI medium. Immune cells (CD4^+^ T cells or Jurkat cells) were first culture in 96 well plates with 50 µg/mL NP administration as mentioned above for 6-48 hours. Then, cells were centrifuged and resuspended in the collagen solution at a density of 1.5×10^5^ cells/mL. A volume of 100 µL per well was seeded into 96-well plates, followed by 20 minutes incubation at 4 ^◦^C to allow cells to settle. Collagen gels were polymerized for 1 hour at 37 ^◦^C and rehydrated with complete RPMI.

Brightfield live-cell imaging of Jurkat and CD4^+^ T cell motility was performed using a Leica Stellaris 5 confocal microscope equipped with a 20×/0.75 NA objective and a humidified chamber (Tokai Hit) maintained at 37 ^◦^C and 5% CO_2_. Images were acquired every 2 minutes for a total duration of 3 hours at 1024×1024 scanning resolution. Individual cell boundaries were segmented from acquired images using CellPose (*59*). Cell centroids were tracked using TrackPy, a Python package based on the Crocker-Grier algorithm, to classify motile trajectories over 30 successive frames (i.e., 1 hour) and quantify 79 single-cell motility features.

### 2D Cell Migration Assay and Data Analysis

Glass-bottom 24-well plates were coated with 50 µg/mL Collagen-I (Enzo) and incubated at 37 ^◦^C for 1 hour. 5,000 3T3-GFP-CaaX cells or 10,000 RAW 264.7 cells per well were seeded and allowed to attach and spread overnight. Prior to imaging, 500 µL of media containing 0, 20, 50, or 200 µg/mL NPs was added to each well and incubated for 6 hours. For RAW 264.7 cells, 20 ng/mL interleukin-6 (IL-6) was co-administered with NPs for 6 hours to stimulate macrophage migration (*60*), followed by staining with CellTracker Green CMFDA fluorescent probe (Invitrogen) at 1:10,000 dilution for 30 minutes, according to the manufacturer’s protocol. All wells were washed three times with DPBS prior to imaging. Live-cell imaging of 3T3-GFP-CaaX and stained RAW 264.7 cells was performed using an epi-fluorescence microscope equipped with a stage-top incubator at 37 ^◦^C and 5% CO_2_. Time-lapse images were acquired every 5 minutes over a 2 hour period. Red fluorescence from NPs was simultaneously imaged to confirm NP association.

Individual cells were tracked using a custom MATLAB script. First, a Gaussian-blurred fluorescence image was thresholded to generate a rough binary cell mask. Each mask was then expanded by 10 pixels (2.27 µm) in all directions to ensure full cropping. A rectangular bounding box was drawn around each cell using its maximum x and y dimensions, and cells with overlapping boxes were excluded from analysis. After segmentation across all frames, a frame-to-frame tracking algorithm was applied based on geometric centroid position. Specifically, each cell at time *t* (*C_t_*) was assigned to its nearest neighbor at *t* + Δ*t* (*C_t_*_+Δ_*_t_*), followed by cross-validation: *C_t_*_+Δ_*_t_* had to identify *C_t_* as its closest cell at frame *t*. Tracks were discarded if cross-validation failed or identified multiple matches, which commonly occurred due to migration, detachment, or overlapping cells. The geometric centroid of each validated cell was then used to construct single-cell migration trajectories.

### Single-Cell Trajectory Dimensionality Reduction and Clustering

All 79 single-cell motility features were standardized using z-score normalization and subsequently log-transformed. Principal Component Analysis (PCA) was applied to identify the minimum number of components that captured at least 75% of the total variance. Uniform Manifold Approximation and Projection (UMAP) was used to embed the standardized features into a two-dimensional Euclidean space, with each point representing a single-cell trajectory. K-means clustering was applied to group trajectories with similar motility patterns. Resulting clusters were labeled in ascending order according to their mean cell speed.

### Fluorescence Recovery After Photobleaching for Actin Dynamics

Fluorescence recovery after photobleaching (FRAP) experiments were performed using a Zeiss LSM800 confocal microscope equipped with a 63x oil-immersion objective and a laser bleaching module. NIH 3T3 cells expressing EGFP-actin were seeded onto collagen-coated glass-bottom 24-well plates under the same conditions used for 2D migration assays. Rectangular regions of interest were manually positioned at the leading edge to target lamellipodial actin networks. Photobleaching was performed using a 488 nm laser at 100% power for 200 iterations. Immediately after bleaching, time-lapse imaging was acquired for 2 minutes at 1-second intervals to monitor fluorescence recovery. The turnover half-time was defined as the time required to reach 50% recovery of the pre-bleach fluorescence intensity after background correction and normalization.

### Preparation of Media with Varying Viscosities and Osmolarities

Media with varying viscosities were prepared as described in Refs. 28,29. In brief, a 3% methylcellulose (65 kDa) stock solution in Iscove’s Modified Dulbecco’s Medium (IMDM; R&D Systems) was diluted with appropriate amounts of cell culture media. Up to 20% of the methylcellulose stock solution was used to generate media with a viscosity of 8 cP. The proportion of IMDM was maintained at 20% in all media, including the 0.8 cP control condition. The osmolarity change due to methylcellulose addition was negligible. Viscosity was measured at 37 ^◦^C using a Cannon-Fenske capillary viscometer.

Media with varying osmolarities were prepared following Refs. 33, 34, while maintaining consistent nutrient concentrations. 80% hypotonic media were prepared by mixing 80% cell culture media with 20% ultra-pure water. 90% hypotonic media were prepared by mixing 80% cell culture media, 10% Dulbecco’s phosphate-buffered saline without calcium and magnesium (DPBS; Sigma-Aldrich), and 10% ultra-pure water. Isotonic media were prepared by mixing 80% cell culture media with 20% DPBS. 110% and 120% hypertonic media were prepared by supplementing with 35 mM and 70 mM D-sorbitol (MilliporeSigma), respectively. Osmolality was measured using a model 3320 osmometer (Advanced Instruments).

### Hydrostatic Pressure Application

A customized air pressure-based incubation system was used to apply hydrostatic pressure. MDCK or HUVEC cells were seeded into 24-well plates as described above, and plates were placed inside an air-tight plastic container. The container was connected to a high-precision digital pressure controller (OB1, Elveflow) and a piezoelectric pressure sensor (MPS, Elveflow) for live pressure monitoring. The pressure controller was connected to compressed air containing 5% CO_2_ and a vacuum pump, enabling precise control of internal pressure to ±3.5 kPa. A flow control valve (McMaster-Carr) was installed on the container to allow limited air exchange. The entire pressure system was placed inside a standard cell culture incubator to maintain temperature at 37 ^◦^C.

### Statistical Analysis

Error bars represent the mean and standard deviation from at least three biological replicates. For single-cell experiments, the number of cells analyzed per condition is provided in the corresponding figure captions. The D’Agostino-Pearson omnibus normality test was used to assess data distribution. For normally distributed data, comparisons between two groups were performed using Student’s *t*-test, and comparisons among more than two groups were conducted using one-way ANOVA followed by Holm-Šídák multiple comparison test. For data normalized to control conditions, paired *t*-tests or repeated-measures (RM) one-way ANOVA was used. For non-normally distributed data, comparisons between two groups were performed using the nonparametric Mann-Whitney *U* -test, and comparisons among more than two groups were conducted using Kruskal-Wallis tests followed by Dunn’s multiple comparison test. All statistical analyses were performed using MATLAB 2021b (MathWorks) or GraphPad Prism 10 (GraphPad Software). Statistical significance was defined as *p <* 0.05. Significance annotations were: ns (*p >* 0.05), * (*p <* 0.05), ** (*p <* 0.01), *** (*p <* 0.001), and **** (*p <* 0.0001).

## Acknowledgments

This work was supported by NIH R01 GM134542 (S.X.S.).

## Supplementary Figures

**Figure S1:**
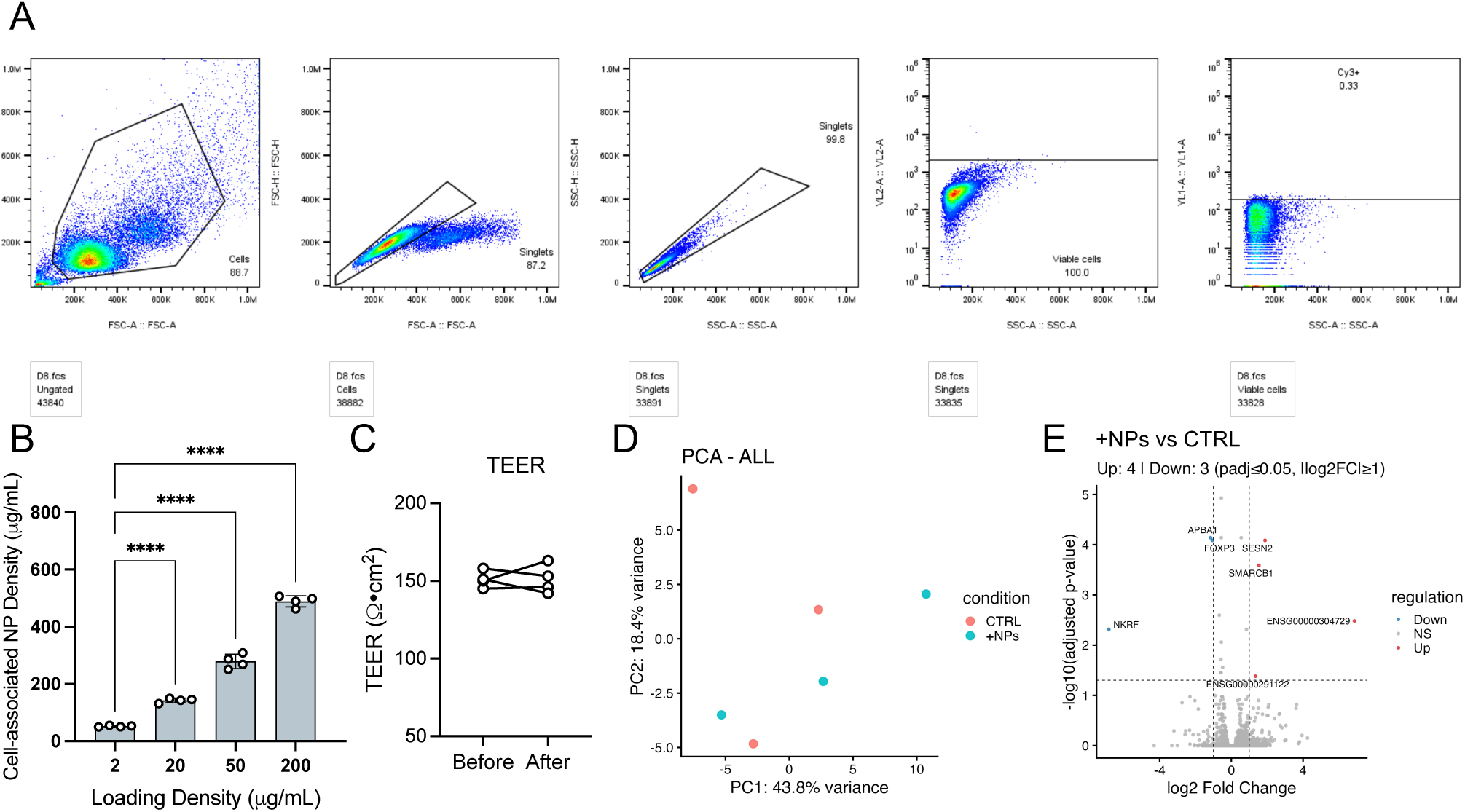
Gating strategy for flow cytometry, NP association, MDCK II confluence measurement, and RNA sequencing. (A) Gating strategy example of NP-association for flow cytometry data analysis. Gating was first based on FSC/SSC together with FSC-A/FSC-H and SSC-A/SSC-H (singlet populations). The live cell population within the gate was further analyzed based on Cy3 signal. (B) Quantification of cell-assocaited NP density at different NP loading times in the MDCK II monolayer. 2, 20, 50, and 200 µg/mL NPs were loaded for 6 hours, followed by washing and epifluorescence imaging. The mass density was then calculated based on the cell height measured in Fig. 1(A) and the calibration curve in 1(C). N = 4. Error bars represent the standard deviation. (C) MDCK II monolayer confluence examined by transepithelial electrical resistance (TEER) measurement before and after adding 20 µg/mL NPs for 6 hours. (D and E) Principal-component analysis (PCA; D) and volcano plots showing differentially expressed genes (DEGs; E) for CD4 T cells RNA expression with or without 50 µg/mL NPs for 48 hours. N = 3.

**Figure S2:**
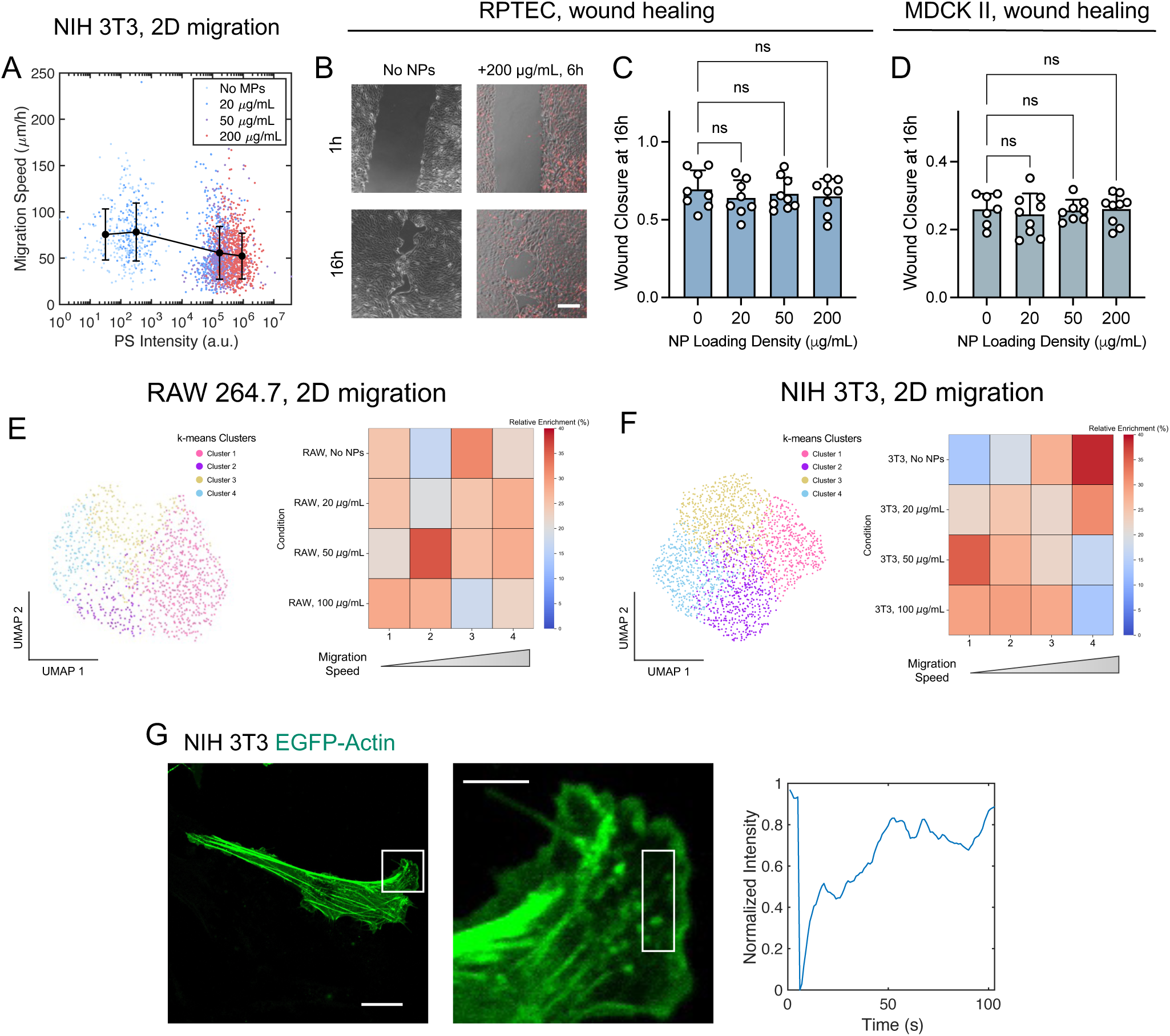
Cell migration analysis, wound healing, and actin turnover dynamics. (A) Single cell migration speed of 3T3 cells as shown in Fig. 3D versus their NP intensity. Error bars represent the mean ± standard deviation of migration speed within intensity bins. (B and C) Representative images (B) and quantification (C) of relative wound closure at 16 hours in RPTEC epithelial monolayer scratch-wound healing assays. N = 8-9. Cells were treated with NPs for 24 hours.(D) Quantification of relative wound closure at 16 hours in MDCK II epithelial monolayer scratch-wound healing assays. N = 7-9. Cells were treated with NPs for 24 hours. (E and F) UMAP of motility parameters (E) and heatmap of relative enrichment in each cluster motility cluster (F) for NIH 3T3 and RAW 264.7 cells under varying NP conditions. (G) Representative images and fluorescent recovery dynamcis of FRAP experiment on EGFP-actin expressing NIH 3T3 cells. Scale bar = 20 µm (left) and 5 µm (right).

**Figure S3:**
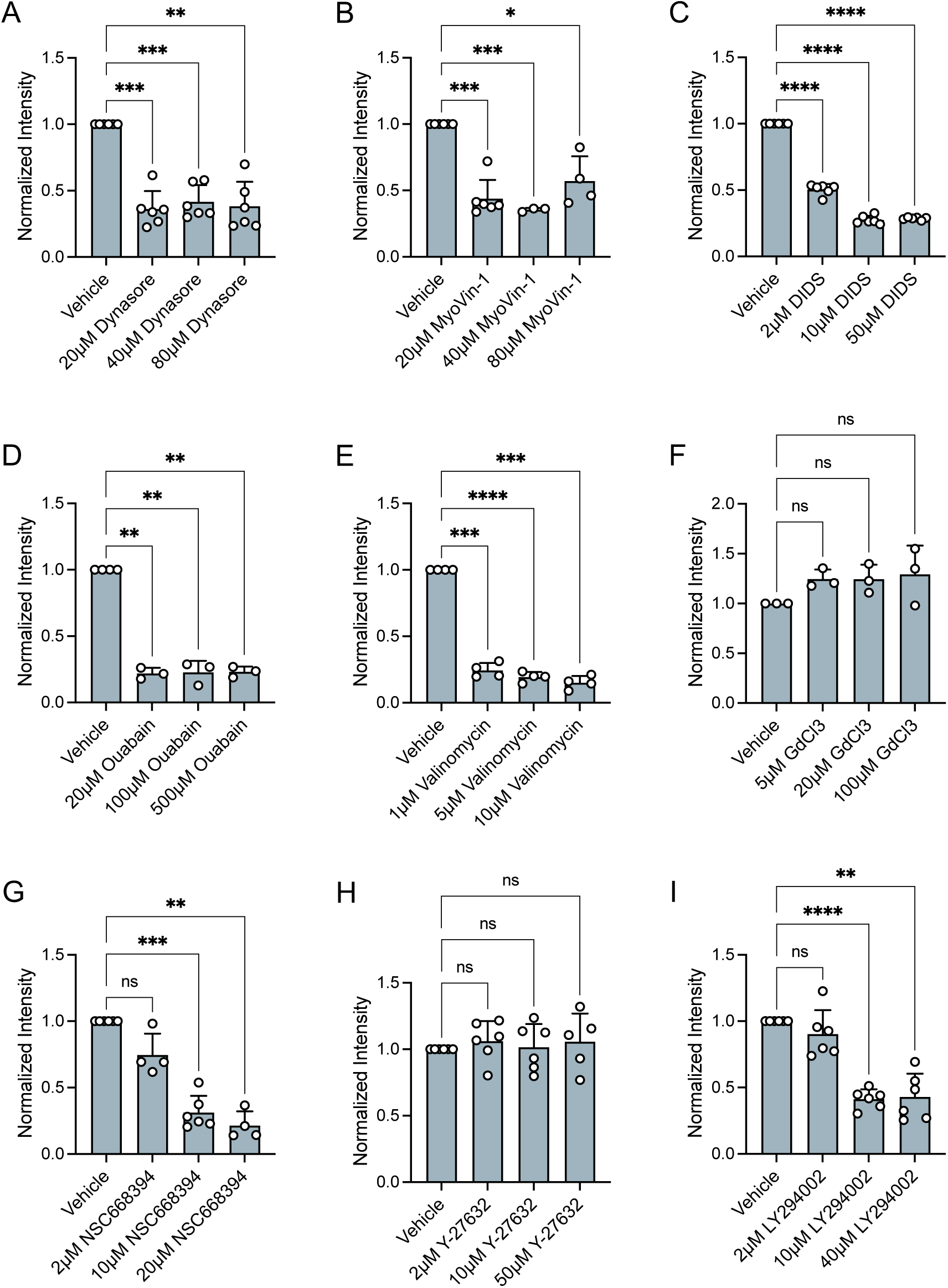
Titrition of inhibitor concentrations on NP-cell association. Dose-dependent response of NP association to MDCK II monolayers under different drug treatment. N = 3-6. Error bars represent standard deviation. Paired *t*-test or RM one-way ANOVA was used.

**Figure S4:**
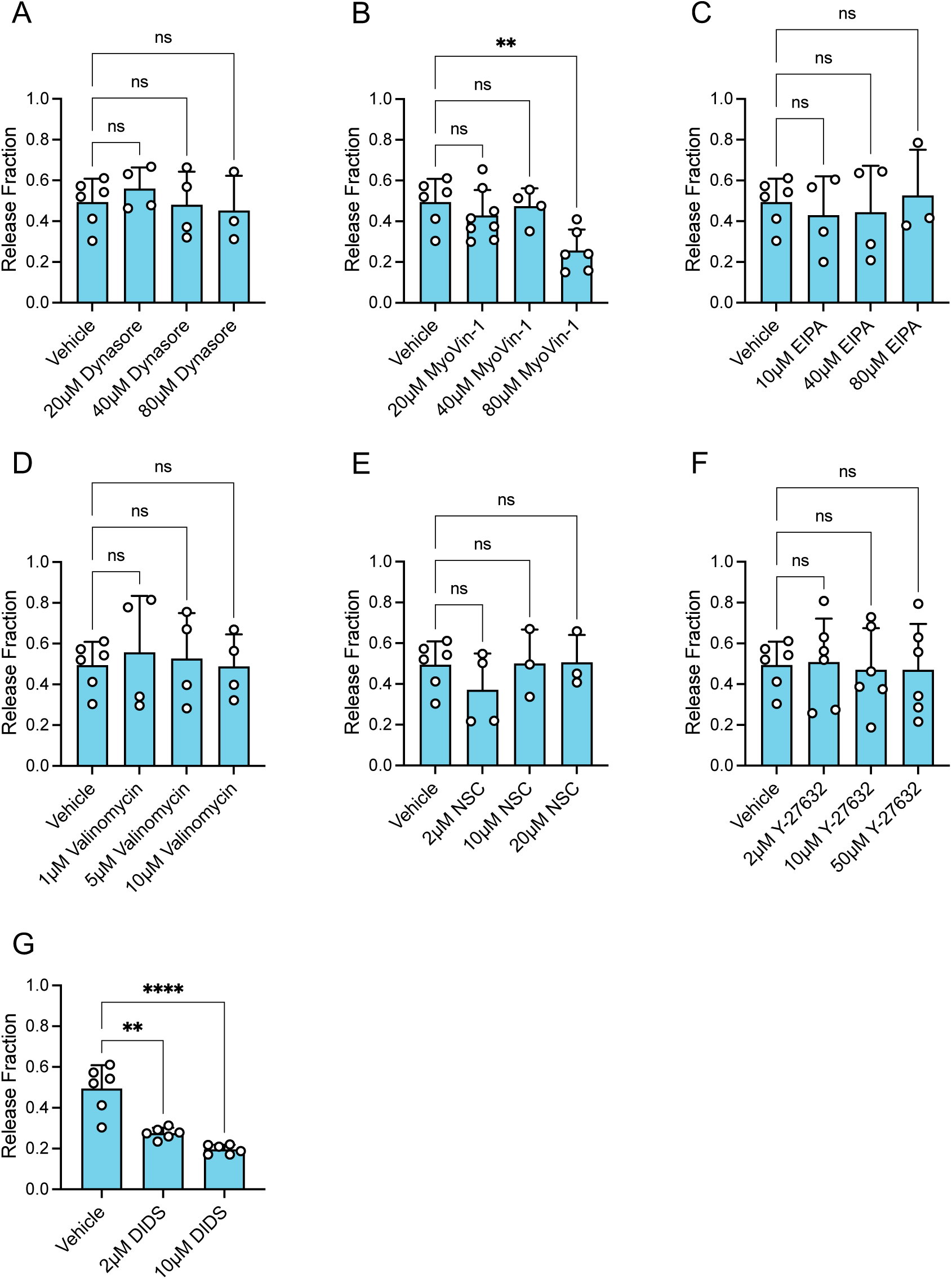
Titrition of inhibitor concentrations on NP release from cells. Dose-dependent response of relative NP release from MDCK II monolayers under various drug treatments. All samples were first incubated with 20 µg/mL NPs for 6 hours without drug treatment, followed by extensive washing and imaging to quantify the initial level of cell-associated NPs. Cells were then incubated in NP-free media containing inhibitors for 24 hours, after which NP fluorescence was measured again. The release fraction was calculated as the relative decrease in fluorescence intensity over this period. Conditions that compromised monolayer integrity due to cytotoxicity were excluded. N = 3-6. Error bars represent standard deviation. Student’s *t*-test or one-way ANOVA was used.

**Figure S5:**
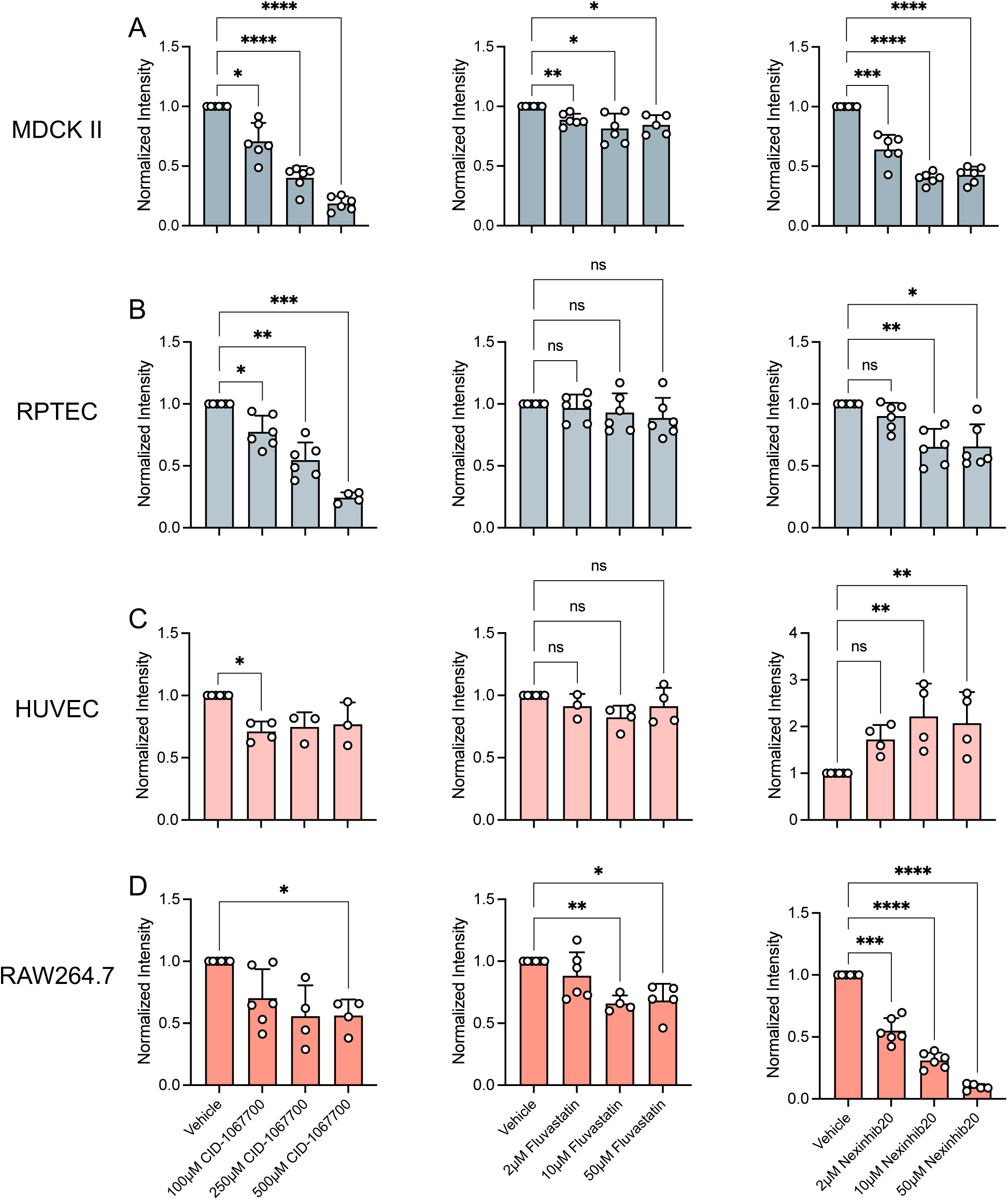
Inhibition of Rab GTPases affects NP-cell association on different cell types. Dose-dependent response of NP association to MDCK II (A), RPTEC (B), HUVEC (C), and RAW264.7 (D) under CID-1067700 (inhibits Rab7), Fluvastatin (inhibits statin), and Nexinhib20 (inhibits Rab27) treatments. N = 4-6. Error bars represent standard deviation. Paired *t*-test or RM one-way ANOVA was used.

**Figure S6:**
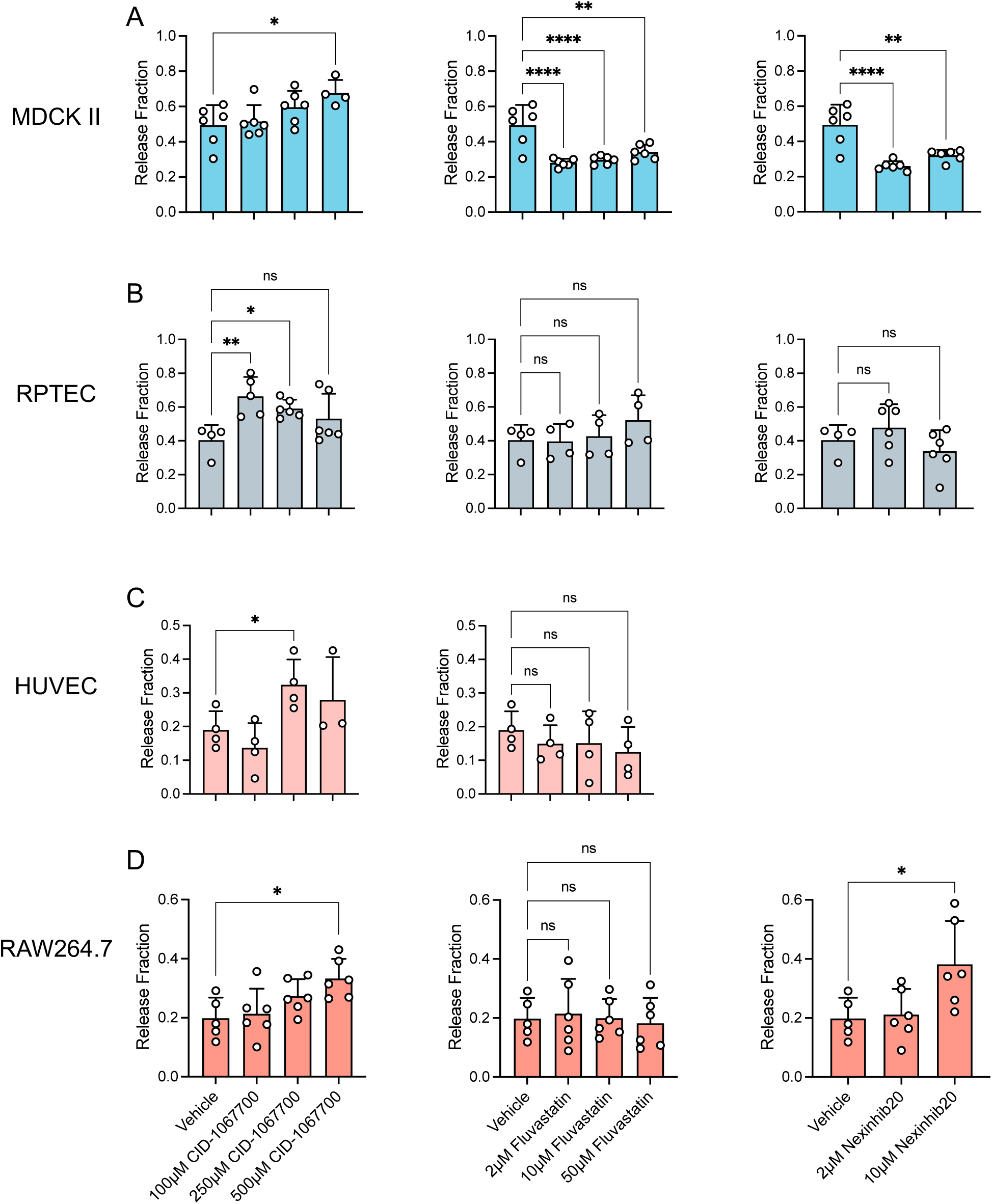
Inhibition of Rab GTPases affects NP release from cells on different cell types. Dose-dependent response of relative NP release from MDCK II (A), RPTEC (B), HUVEC (C), and RAW264.7 (D) under CID-1067700 (inhibits Rab7), Fluvastatin (inhibits statin), and Nexinhib20 (inhibits Rab27) treatments. Conditions that compromised monolayer integrity or reduced viability due to cytotoxicity were excluded. N = 3-6. Error bars represent standard deviation. Student’s *t*-test or one-way ANOVA was used.

**Figure S7:**
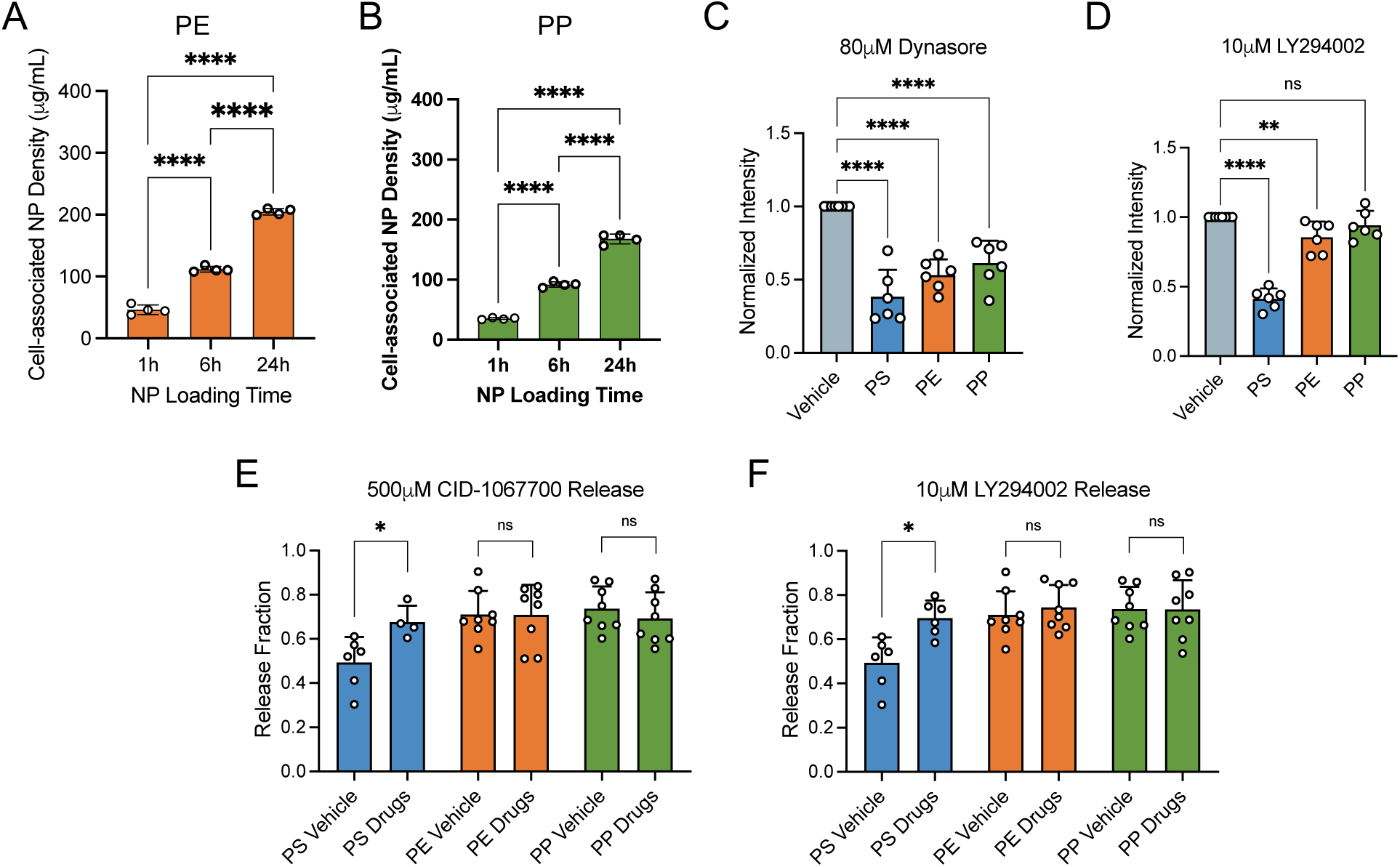
Association of PE and PP NPs. (A and B) Quantification of cell-associated PE (A) and PP (B) NP density at different NP loading times in the MDCK II monolayer. 20 µg/mL NPs were loaded for 1, 6, and 24 hours, followed by washing and epifluorescence imaging. (C and D) PS, PE, and PP NP association in MDCK II monolayers treated with dynasore (C) or LY294002 (D). N = 6. (E and F) Release fraction of PS, PE, and PP NPs in MDCK II monolayers treated with CID-1067700 (E) and LY294002 (F). N = 4-8. (A-F) Error bars represent standard deviation. One-way ANOVA or Student’s *t*-test was used. RM one-way ANOVA was used in (C and D).

**Figure S8:**
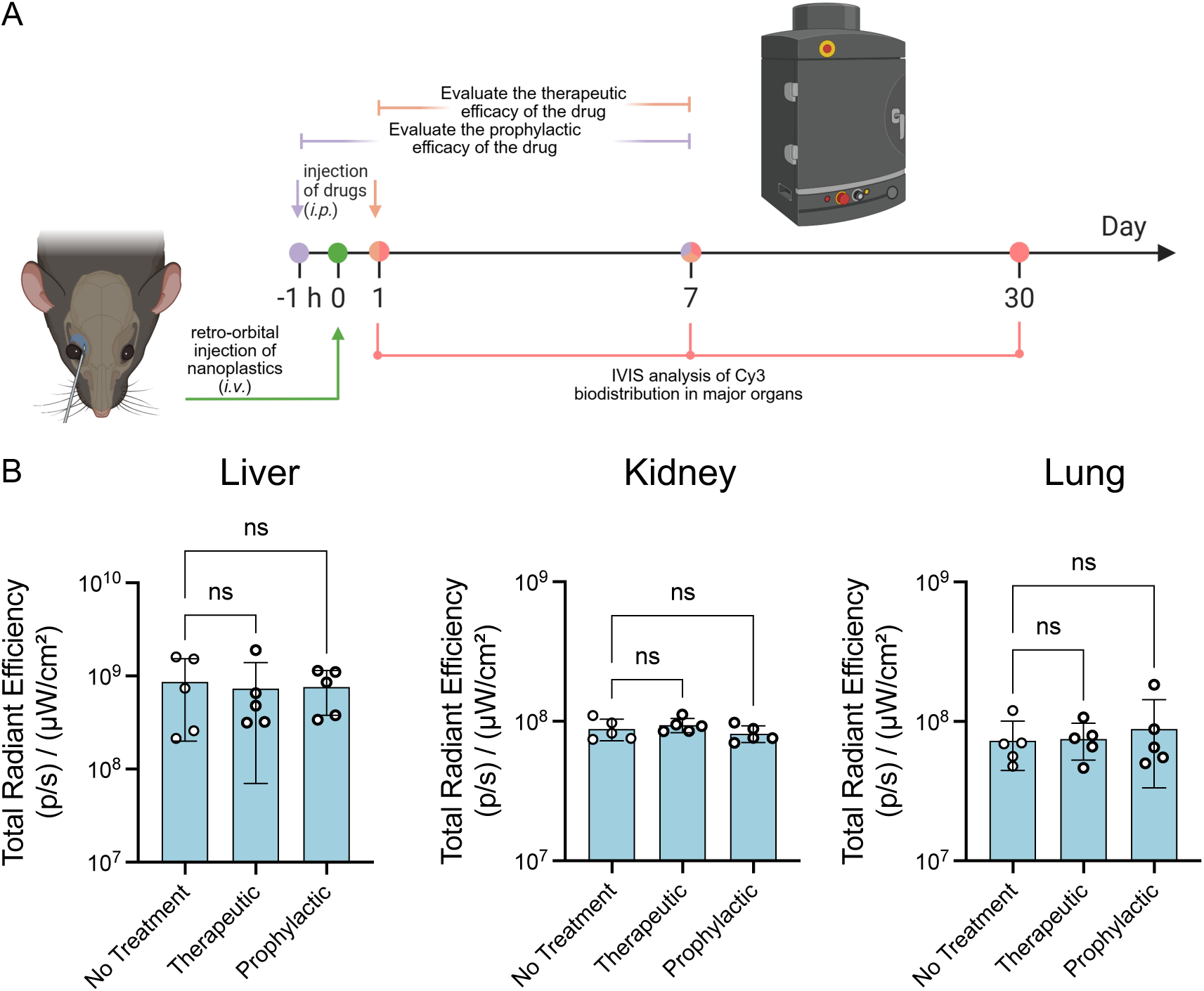
Therapeutic and prophylactic efficacy of CID-1067700 on NP biodistribution in mice. (A) Schematic of the therapeutic and prophylactic treatment schemes. CID-1067700 (16 mg/kg, dissolved in 50 µL DMSO) was administered by intraperitoneal (i.p.) injection either 1 hour before NP administration (prophylactic assessment) or 1 day after NP administration (therapeutic assessment), following the NP injection protocol shown in Fig. 1F. Mice were sacrificed 7 days post-injection, and the biodistrubiton of NPs in major organs was assessed using IVIS. (B) Total radiant efficiency of NPs in liver, kidney, and lung with or without CID-1067700 treatment. Error bars represent standard deviation. One-way ANOVA was used.

## Notes

### Competing Interest Statement

The authors have declared no competing interest.

